# Comparative landscape of extracellular vesicles across three pathogenic trematodes identifies a key conserved miRNA cargo involved in host liver fibrosis

**DOI:** 10.64898/2026.11.19.767961

**Authors:** Lu Liu, Zhigang Lu, Shiyu Zhu, Chuantao Fang, Xiaoli Yan, Chunren Wang, Yanyan Zhang, Feng Liu, Maria Pakharukova, Guofeng Cheng

**Affiliations:** Shanghai Tenth People’s Hospital, Institute for Infectious Diseases and Vaccine Development, Tongji University School of Medicine, Shanghai, China; Clinical Center for Brain and Spinal Cord Research, Tongji University, Shanghai, China; Affiliated Shanghai Blue Cross Brain hospital, Shanghai 200020, China; Faculty of Natural Sciences, University of Hohenheim, Stuttgart, Germany; Teaching Laboratory Center, Tongji University School of Medicine, Shanghai, China; Key Laboratory of Bovine Disease Control in Northeast China, Ministry of Agriculture and Rural affairs, College of Animal Science and Veterinary Medicine, Heilongjiang Bayi Agricultural University, Daqing 163319, Heilongjiang Province, PR China; Institute of Cytology and Genetics, Siberian Branch of Russian Academy of Sciences (ICG SB RAS), Novosibirsk, Russia

**Keywords:** Trematodes, Extracellular vesicles, miRNA, Liver fibrosis

## Abstract

Liver fibrosis caused by *Schistosoma japonicum*, *Clonorchis sinensis*, and *Opisthorchis felineus* represents a major global public health burden. Here, we systematically characterized extracellular vesicles (EVs) released by key stages of these parasites and uncovered, for the first time, a conserved repertoire of EV-derived miRNAs shared across all three species. By integrating animal models, single-cell RNA sequencing, and bioinformatic analyses, we comprehensively identified the targetome of *miR-36-3p*, a cross-species conserved miRNA cargo. Mechanistically, we found that *miR-36-3p* directly targets *Atox1* in Kupffer cells, driving intracellular copper and reactive oxygen species (ROS) accumulation and thereby activating hepatic stellate cells. Strikingly, *in vivo* delivery of an AAV8-encoded *miR-36-3p* sponge markedly attenuated liver fibrosis across all three infection models. Collectively, these findings establish a conserved, parasite-derived EV miRNA cargo as a previously unrecognized driver of host liver pathology and identify it as a promising therapeutic target for preventing parasite-induced liver fibrosis.

## 1 Introduction

Liver fibrosis caused by zoonotic trematodes such as *Schistosoma japonicum* (*S. japonicum*), *Clonorchis sinensis* (*C. sinensis*), and *Opisthorchis felineus* (*O. felineus*) remains a substantial public health problem ^1, 2, 3^. Currently, at least 286.2 million people are at risk of infection with these trematodes ^4, 5, 6^. More worryingly, these infections can result in chronic hepatobiliary inflammation, fibrosis, and even an elevated cancer risk ^5^.

Long-term studies indicate that the primary etiology of the pathogenesis caused by these trematodes lies in the excretory/secretory products (ESPs) released during key developmental stages ^7^. Given that extracellular vesicles (EVs) secreted by almost all cells are recognized as important regulators in many physiological and pathological processes ^8^, several studies have focused on the functions of EVs from these trematodes including *S. japonicum* eggs, adult *C. sinensis* or *O. felineus* ^9, 10, 11^. Unfortunately, these studies either focused on a single parasite or were restricted to basic EV characterization and preliminary functional analyses of the EV-derived miRNA cargo. For instance, in *S. japonicum* eggs, parasite EVs were found to contain not only small RNAs but also miRNAs such as *miR-33* and *miR-71a* that may regulate liver fibrosis ^12, 13^. *C. sinensis* EVs were found to foster the malignant progression of cholangiocarcinoma, and their miRNA cargo, such as *Csi-let-7a-5p*, triggers pro-inflammatory macrophage activation, leading to biliary injury ^14^. In *O. felineus*, EV-like vesicles are secreted and subsequently taken up by human cholangiocytes ^9^. However, it remains unknown whether these related trematodes, which induce similar clinical pathologies in final hosts after infection, share a common molecular basis and regulatory mechanism driving host liver fibrosis.

In this study, we systematically profiled the landscape of EVs secreted during the key pathogenic stages of these three trematodes. Notably, we identified a conserved repertoire of EV miRNA cargos shared across these trematode pathogens. By integrating animal models with single-cell RNA sequencing and bioinformatics, we comprehensively characterized the targetome of *miR-36-3p*, a common EV miRNA cargo shared by all three parasites. Mechanistically, we demonstrate that *miR-36-3p* targets *Atox1* in Kupffer cells, thereby elevating copper and ROS levels and consequently regulating HSC activation. Moreover, *in vivo* administration of an AAV8-delivered miRNA sponge markedly attenuated liver fibrosis, demonstrating that *miR-36-3p* not only acts as a pathogenic mediator derived from parasite EVs but also serve as a promising therapeutic target for liver fibrosis.

## 2 Method and Materials

### 2.1 Ethical statement

Animal protocols in the present study were approved by the Animal Ethics Committee of Shanghai 10^th^ People Hospital of Tongji University (ID number: SHDSYY-2023-P0036).

### 2.2 Animal models

Animal models for *S. japonicum* infection were established as described previously ^15^. Male New Zealand White rabbits (3–6 months old) and mice were percutaneously challenged with certain number of cercariae^11^, which were shed from positive *Oncomelania hupensis* snails obtained from Jiangxi Provincial Institute of Parasitic Diseases (Jiangxi, China). Rabbits/hamsters were used for *C. sinensis* and *O. felineus* infection ^16, 17, 18^. All animals were housed under standard conditions with free access to food and water. Parasites were harvested at 56 days post-infection (dpi).

### 2.3 Eggs and adult worm collection

*S. japonicum* eggs were obtained as previously described ^19, 20^. Briefly, following euthanasia, rabbit livers were perfused with PBS to remove residual blood. The tissue was homogenized in pre-cooled 1.2% NaCl, and the resulting suspension was sequentially filtered through 160-μm and 360-μm meshes. The material retained on the 360-μm mesh, which contained the eggs, was collected and transferred to a beaker. The eggs were further washed three times with 1.2% NaCl at 4°C via gravity sedimentation. The purified eggs were collected and washed three times with PBS.

### 2.4 EVs collection

EVs were isolated from the egg/worm culture condition medium following a previously established protocol^21^. Briefly, *Schistosoma japonicum* eggs, *Clonorchis sinensis* worms, and *Opisthorchis felineus* worm as well as the blank (without worms/eggs) were cultured in the RPMI 1640 with 10% FBS with EVs free and 1% penicillin-streptomycin (BBI, Shanghai, China), and every 8 hours changed medium. Dead cells and cell debris were pelleted by centrifugation at 1000 × g for 10 min and then 10000 × g for 30 min to remove and the supernatant was filtrated through 0.22 μm membrane. The filtrate was then ultracentrifuged at 100,000 × g for 2 h at 4℃. The supernatant was collected for the usage of control. The EV pellet was resuspended in PBS. In parallel, the EV depleted supernatant was used for control. The condition medium without worms/eggs were also subject to EV process for experiment control. The endotoxin assay for each isolated EVs was performed as described previously^22^.

### 2.5 Western blotting

Antibodies for CD63, CD81, HSP 70, and TSG101 were used to analyze the isolated EVs by Western blotting ^23^. Primary antibodies against calnexin (1:5000 dilution, Abcam, Cambridge, UK, Catalog# ab275018) and BSA (1:5000 dilution, Sangon, Shanghai, China; Catalog# D220272) were also applied. The secondary antibodies were horseradish peroxidase (HRP)-conjugated secondary antibodies (1:5000 dilution, Yasen, Shanghai, China; Catalog# 33101ES). Protein bands were finally visualized using an enhanced chemiluminescence (ECL) (Beyotime Biotechnology, Shanghai, China).

### 2.6 Transmission electron microscopy

EV morphology was characterized using a JEOL 1200EX II transmission electron microscope (JEOL, Tokyo, Japan) equipped with a Gatan digital camera (Gatan, Pleasanton, USA).

### 2.7 Nanoparticle tracking analysis

Nanoparticle tracking analysis (NTA) was conducted using a NanoSight NS300 system (Malvern Instruments, Malvern, UK). EVs were mixed with PBS and loaded into the sample chamber. Three independent measurements were taken for each sample ^23^.

### 2.8 Endotoxin contamination

Endotoxin contamination of the samples was assessed with the ToxinSensor™ Chromogenic LAL Endotoxin Assay Kit (GenScript, Nanjing, China) following the supplier’s protocol with minor modifications^22^. The assay is a quantitative endpoint chromogenic LAL method. In brief, 0.1 mL of sample or *E. coli* endotoxin standard was mixed with 0.1 mL of reconstituted LAL reagent and incubated at 37 °C for 60 min. After adding 0.1 mL of chromogenic substrate and a further 6 min at 37 °C, the reaction was terminated and stabilized, and absorbance was read at 545 nm on a SPECTRA MAX 190 spectrophotometer. Endotoxin concentrations were determined from a standard curve prepared with the *E. coli* endotoxin standards.

### 2.9 RNA sequencing and bioinformatics analysis

Small RNA sequencing was carried out at the BGI (Shenzhen, China) using an Illumina platform. The following bioinformatic analysis was conducted as described previously ^14^.

### 2.10 Total RNA isolation and cDNA synthesis

Total RNA was extracted from cells or livers using TRIzol™ Reagent (Thermo Fisher Scientific, USA) and RNA concentrations were assessed with a NanoDrop™ One spectrophotometer (Thermo Fisher Scientific, USA). cDNA was synthesized using the PrimeScript™ RT reagent Kit (Takara Bio, Kusatsu, Japan). For miRNAs, cDNA was synthesized using the miRNA 1st Strand cDNA Synthesis Kit (Vazyme, Nanjing, China) with stem-loop RT primers (2 μM), 10× RT mix (2 μL), and HiScript II Enzyme Mix (2 μL) in a 20 μL reaction. The reaction was incubated at 25 °C (5 min), 45℃ (15 min), and 85 °C (5 min) for enzyme inactivation. The synthesized cDNA was diluted and stored at −20 °C for the following analyses.

### 2.11 Cell culture

RAW264.7 cells, JS1 cells (provided by Prof. Pan’s laboratory at the Naval Medical University), and Kupffer cells (ImKC) were cultured in high glucose DMEM medium (BBI, Shanghai, China) with the addition of 1% (v/v) penicillin-streptomycin (BBI, Shanghai, China) and 10% fetal bovine serum (Thermo Fisher Scientific, USA).

For the co-culture experiment, JS1 cells were seeded into the lower chambers of 12-well Transwell plates (Vazyme, Nanjing, China). RAW264.7 cells were treated with parasite EVs or control CCM at a concentration of 20 μg/mL for indicated times. After 24 hours of co-culture, JS1 cells were collected for subsequent analyses. The co-culture assays were conducted utilizing Vazyme Transwell inserts. The upper chambers were seeded with 100 µL of a cell suspension containing 1 × 10⁴ Kupffer cells. After 24 h of stimulation with EVs or mimics, the Transwell inserts were transferred to wells containing JS1 cell cultures.

### 2.12 Primary cell isolation from livers

Liver cells were isolated by *in vivo* perfusion and digestion, as previously described ^24, 25^. Initially, the livers were perfused with HBSS for approximately 5 minutes. This was followed by a 5-minute perfusion with collagenase IV (3.7 U/mouse) (Thermo Fisher Scientific, USA; Catalog# 17104019) and Pronase E (14 mg/mouse) (Coolaber, Beijing, China, Catalog# CP9141). Subsequently, the livers were excised, minced, and incubated for 20 min at 37°C with agitation at 100 rpm in a digestion cocktail comprising Pronase E (25 mg/mouse), collagenase IV (4.4 U/mouse), and DNase I (10 U/mL) (Roche, Basel, Switzerland; Catalog# 04716728001). Cell suspensions were sequentially passed through 70-µm and 40-µm mesh filters, with all subsequent steps performed at 4°C. To eliminate hepatocytes, the samples underwent two consecutive centrifugation steps at 50 × g for 3 minutes each. The remaining liver cells were centrifuged at 800 × g for 7-8 min. Following the removal of the supernatant, red blood cell lysis buffer was used to resuspend and the solution was incubated for 5 min on ice. The non-parenchymal cell (NPC) fraction was layered onto a 15% (w/v) iodixanol (Abcam, Cambridge, UK) solution and centrifuged at 1,280 × g for 20 minutes at 4°C. The intermediate layer, containing Kupffer cells, was carefully collected into a 15-mL centrifuge tube, diluted with DMEM, and centrifuged at 700 × g for 8 minutes to remove residual iodixanol. The cell pellet was then resuspended in DMEM with 10% FBS for subsequent experiments ^26^.

### 2.13 Cell Counting Kit-8 (CCK-8) assay

Cell viability was evaluated using the Cell Counting Kit-8 (Yeasen, Shanghai, China) following manual instruction ^27^. Briefly, cells were plated in 96-well flat-bottom plates (1× 10³ cells/well) and exposed to parasite EVs, EV-depleted supernatant, or transfected *miR-36-3p* mimics and/or antisense. Following the addition of 10 μL CCK-8 reagent to each well, the plates were further incubated at 37°C for 1h. Absorbance was measured at 450 nm on a microplate reader at indicated time points.

### 2.14 Cell migration experiment

Cell migration experiment was performed as described previously ^28^, with slight modifications. JS1 cells were plated in 6-well plates. A linear scratch was introduced in the monolayer cells. Detached cells were removed by washing, and the remaining cells were co-cultured with KCs in a co-culture system for 24 h. Cell migration was observed using an inverted microscope (Motic, Xiamen, China) and quantified by ImageJ software (National Institutes of Health, USA).

### 2.15 EVs labelling with PKH26 and cell uptake

PKH26 labeling of EVs and EV-depleted supernatant (EVDS) were performed as described previously ^29, 30^. The PKH26 dye (Duolaimi, Wuhan, China) was vortexed and centrifuged, and 5 μL dye was added to 50 μL reaction buffer and mixed until dissolved. Then, 50 μL of EVs, EVDS, or control CCM was mixed with 60 μL of dye solution, followed by incubation at 37 °C for 30 min under dark. Quenching was performed using 15 mL sterile PBS (0.22 μm filter) and then ultracentrifuged at 100,000 × g at 4 °C for 2 h and further resuspended in 200 μL sterile PBS. Cells were seeded and allowed to adhere, after which the culture medium was replaced with fresh medium containing PKH26-labelled EVs. Cell internalization was observed using an inverted fluorescence microscopy (LSM 900, ZEISS, Germany).

### 2.16 ALT, AST and HYP assay

The levels of liver hydroxyproline (HYP) were quantified using the Hydroxyproline Assay Kit (Nanjing Jiancheng Bioengineering Institute, Nanjing, China) following the manufacturer’s instructions with absorbance recorded at 550 nm on a microplate reader (Molecular Devices, San Jose, USA). Serum aminotransferase (ALT) and aspartate aminotransferase (AST) activities were determined via the Reitman-Frankel colorimetric assay (Nanjing Jiancheng Bioengineering Institute, Nanjing, China) according to the production manual and absorbance was measured at 505 nm using the same microplate reader.

### 2.17 Immunohistochemistry

Liver tissues were fixed in 4% formalin and embedded in paraffin. Following dewaxing, the sections were incubated overnight at 4°C with primary antibodies against α-SMA (1:500 dilution, Abcam, USA; Catalog #: ab7817) and anti-Atox1 (1:200 dilution, Sangon, Shanghai, China; Catalog#: D123073). Goat anti-rabbit horseradish peroxidase (HRP)-conjugated secondary antibodies (1:500 dilution) (Yasen, Shanghai, China; Catalog# 33101ES) was used as secondary antibodies. The signals were then developed with DAB substrate (Servicebio, Wuhan, China). Representative images were captured using a microscopy (Motic, Xiamen, China).

### 2.18 Histopathological analysis

The histopathological analyses were conducted as described previously ^26, 31^. After staining of liver tissue sections with H&E, the granulomas were quantified using Image J. For the Masson’s staining, the sections were initially stained with Weigert’s iron hematoxylin solution (Yeasen, Shanghai, China) and then stained with Ponceau acid fuchsin solution. After a brief rinse, the sections were differentiated with 1% HCL for 2 seconds and blue with PBS for 8 minutes and rinsed again. Finally, the sections were stained with aniline blue solution for 15minutes. The sections were then mounted with neutral gum and examined and imaged using an optical microscope (Motic, Xiamen, China).

### 2.19 Immunofluorescent staining

The slides with cells were fixed in 4% paraformaldehyde (Servicebio, Wuhan, China) and then treated with 0.1% Triton X-100 (Applygen, Beijing, China). Then, the cells were incubated with 5% BSA (Yeasen, Shanghai, China). The primary antibodies including anti-iNOS and anti-CD86 (both at 1:2000 dilutions; Abcam, Cambridge, UK, Catalog#: ab283655 and ab239075), as well as anti-α-SMA and anti-Col I / III (1:500 dilutions, Abcam, Cambridge, USA; Catalog# ab7817) were used. The secondary antibodies were horseradish peroxidase (HRP)-conjugated secondary antibodies (1:5000 dilution, Yasen, Shanghai, China; Catlog# 33101ES). Subsequently, the signals were then developed with CY5 or CY3 substrates for fluorescence amplification (Yeasen, Shanghai, China) was incubated for 15 min at room temperature. The target areas were quantified using Image J.

### 2.20 Cell transfection

Kupffer cells were placed in either 12-well plates. Then, the cells were transfected in a serum-free medium with miRNA mimics or siRNA-Lipo3000 complexes (Vazyme, Nanjing, China) at a concentration of 80 nM. After 8 h of transfection, the medium was changed, and then added DMEM with 10% fetal bovine serum (FBS) (Vazyme, Nanjing, China) and 1% penicillin–streptomycin (BBI, Shanghai, China). The cells were either collected or given more treatments as indicated time ^32^. The sequences of *miR-36-3p* mimics, and *Atox1* siRNA are listed in supplementary Table 4.

### 2.21 Liver cell isolation for scRNA-seq analyses

Liver cells were isolated as previously described ^32^. Briefly, the murine liver was perfused with calcium-free HBSS, containing 0.2 mg/mL EDTA. After sequential perfusion with 0.4 mg/mL pronase E (Coolaber, Beijing, China; Catalog# CP9141) and 0.2% collagenase type IV (Thermo Fisher Scientific, USA; Catalog# CP9141), liver was further digested in HBSS containing 0.2% collagenase type IV, 0.4 mg/mL pronase E, and 0.1 mg/mL DNase I (Roche, Basel, Switzerland, Catalog# CP9141). After adding 10 mL of cold DMEM, the digestion was terminated. The cell suspension was spun down by centrifugation (50 × g, 3 min), and the resulting filtrate was sequentially passed through 70-μm and 40-μm nylon cell strainer (Yeasen, Nanjing, China). The flow was then treated with red blood cell lysis buffer ((BioLegend, San Diego, USA) for 5 min in ice. The cell suspension was subjected to density gradient centrifugation using 20% Optiprep (Abcam, Cambridge, UK) to remove dead cells. Cell viability was determined by trypan blue exclusion. The purified NPCs were then subjected to scRNA-seq droplet preparation and library construction using the MGI DNBelab C4 Single-Cell RNA library Prep Set at BGI (Shenzhen, China).

### 2.22 Single-cell RNA data processing, integration, clustering and differential expression

Raw sequencing data from both cDNA and Oligo libraries were concurrently processed using the dnbc4tools pipeline (version 2.1.3). Briefly, dnbc4tools extracted cell barcodes and UMIs from the Oligo library reads and matched them with the transcript reads from the cDNA library. Clean cDNA reads were aligned to the reference genome (GRCm39) using the integrated STAR aligner and annotated to functional genomic regions. Automated bead merging and cell calling algorithms were applied to distinguish true single cells from ambient background noise. The filtered data matrix from dnbc4tools was imported into Seurat 5.1.0 and constructed to a Seurat Object with min.cells=3, min.features=200. The data were subjected to doublet detection using scDblFinder 1.16 and only singlets were kept. To filter low-quality cells, only those with 500-6000 genes, 1000-60000 counts, and less than 5% mitochondrial gene expression were retained. The two data layers (antisense and control) were then merged and subjected to normalization, calculation of high variable genes, scaling and PCA dimensionality reduction. The layers were then integrated using the Harmony Integration method and processed to clustering using dims=1:50 and a resolution of 0.8. Marker genes were calculated using the FindAllMarkers function with the roc method and filtered with AUC > 0.7, pct.1 > 0.7 & pct.2 < 0.5. Differentially expressed genes between antisense and NC cells were calculated using Find Markers function with the default wilcox method using the parameter logfc. Threshold = 0.1, min.pct = 0.1, and then filtered with pval adj < 0.05 & abs(avg_log2FC) > 0.25. This threshold was applied to all cells as well as cells from each cluster.

### 2.23 Bulk RNA-seq data analysis

Sequence reads were aligned to the mouse reference genome and quantified at the gene level. Secondary analysis was performed with iDEP v2.4.4 (https://bioinformatics.sdstate.edu/idep20/). Lowly expressed genes were removed by filtering out reads below 0.5 CPM in at least one sample. Data transformation was conducted using variance-stabilizing transformation (VST), and missing values were imputed by gene median for visualization and clustering. Differential expression analysis was performed with DESeq2 (v1.48.2) using a design of ∼groups, the Wald test, and independent filtering with alpha = 0.1. Genes were considered differentially expressed at Benjamini-Hochberg adjusted p < 0.05 with a fold-change of at least 1.5

### 2.24 Target prediction and bioinformatics analyses

Putative miRNA target genes were computationally predicted using RNAhybrid^33^.To obtain highly reliable target genes, we retained only those genes that were predicted by RNAhybrid and additionally met the following expression criteria: downregulated in KCs following *miR-36-3p* mimics treatment, and upregulated in single-cell RNA-seq data.

### 2.25 Recombinant Adeno-associated Virus (rAAV)

Recombinant adeno-associated virus (rAAV) vectors were purchased from GenePharma Biotechnology (Shanghai, China). The recombinant AAV8 vectors employed in this study were generated, purified, and titrated by GenePharma. For *in vivo* transduction, mice received tail vein injections of 1 × 10¹¹ vector genomes (v.g.) per mouse at 10 days post-infection with *S. japonicum*, *C. sinensis*, and *O. felineus*, as previously reported ^34^.

### 2.26 Colorimetric Assay for Intracellular Copper (Cu²⁺) Measurement

The measurement was conducted using a commercial colorimetric assay kit (Elabscience, Wuhan, China). Briefly, the collected cultured cells were washed three times with ice-cold PBS. Then, the cells were lysed in the lysis buffer on ice for 10 minutes to release their contents and then were centrifuged at 12,000 × g for 10 minutes at 4°C. The supernatants were subject to measure Cu²⁺ concentration. The copper standard from the kit was diluted to create a standard curve from 0.18 to 5 μmol/L. Equal amounts of standards and sample supernatants were mixed with the chromogenic reagent (cupric ion chelator) and incubated at 37°C for 5 minutes. The optical density (OD) of each reaction was measured at 580 nm using a microplate reader (Winooski, Vermont, USA). Copper concentrations in the samples were calculated by comparing them to the standard curve and then normalized with protein concentration (Beyotime, Shanghai, China).

### 2.27 Measurement of intracellular ROS

Intracellular ROS levels were measured using the Reactive Oxygen Species Assay Kit (Beyotime, Shanghai, China) according to the manufacturer’s instructions. Briefly, cells were incubated with 10 µM DCFH-DA at 37°C for 20 min in the dark. After washing three times with PBS to remove residual probe. Subsequently, cells were fixed with 4% paraformaldehyde at room temperature for 10 min and mounted with an anti-fade mounting medium. Fluorescence was detected using an inverted fluorescence microscopy at 488 nm (LSM 900, ZEISS, Germany).

### 2.28 Statistical analyses

Statistical analysis of the differences was conducted Student’s *t* test or one-way ANOVA. Data analyses were conducted using GraphPad Prism software (version 10.1).

## 3 Results

### 3.1 Characterization of isolated extracellular vesicles from key stages of three trematodes

Several animal models infected with the three trematodes were established (**Figure 1A)**. Upon confirmation of bona fide liver pathology in the infected animals (**Figure 1B)**, we isolated the pathogens responsible for hepatic fibrosis including *S. japonicum* eggs, adult *C. sinensis* and adult *O. felineus*, respectively. Subsequently, extracellular vesicles (EVs) from these key stages of trematodes were isolated (**Figure 1A**). Transmission electron microscopy (TEM) analysis indicated that the isolated EVs displayed a typical cup-shaped morphology (**Figure 1C**). Nanoparticle tracking analysis (NTA) showed that the sizes of the isolated EVs ranged from 50 to 200 nm (**Figure 1D**), with corresponding particle concentrations varying from 1.03 × 10¹¹ to 1.44 × 10¹¹ particles/mL. Western blotting analyses indicated that these EVs presented canonical EV marker proteins, including CD63, HSP70, CD81, and Tsg101, whereas the negative control protein, calnexin, was absent (**Figure 1E**). In addition, fetal bovine serum (FBS) was not detected in any of the isolated EV preparations using specific antibodies (**Figure S1**). Moreover, the isolated trematode EVs were not contaminated with endotoxin (**Figure 1F**). Collectively, these characterization data confirm the high purity and integrity of the isolated trematode EVs, providing a reliable foundation for downstream small RNA profiling and functional characterization.

**Figure 1.**
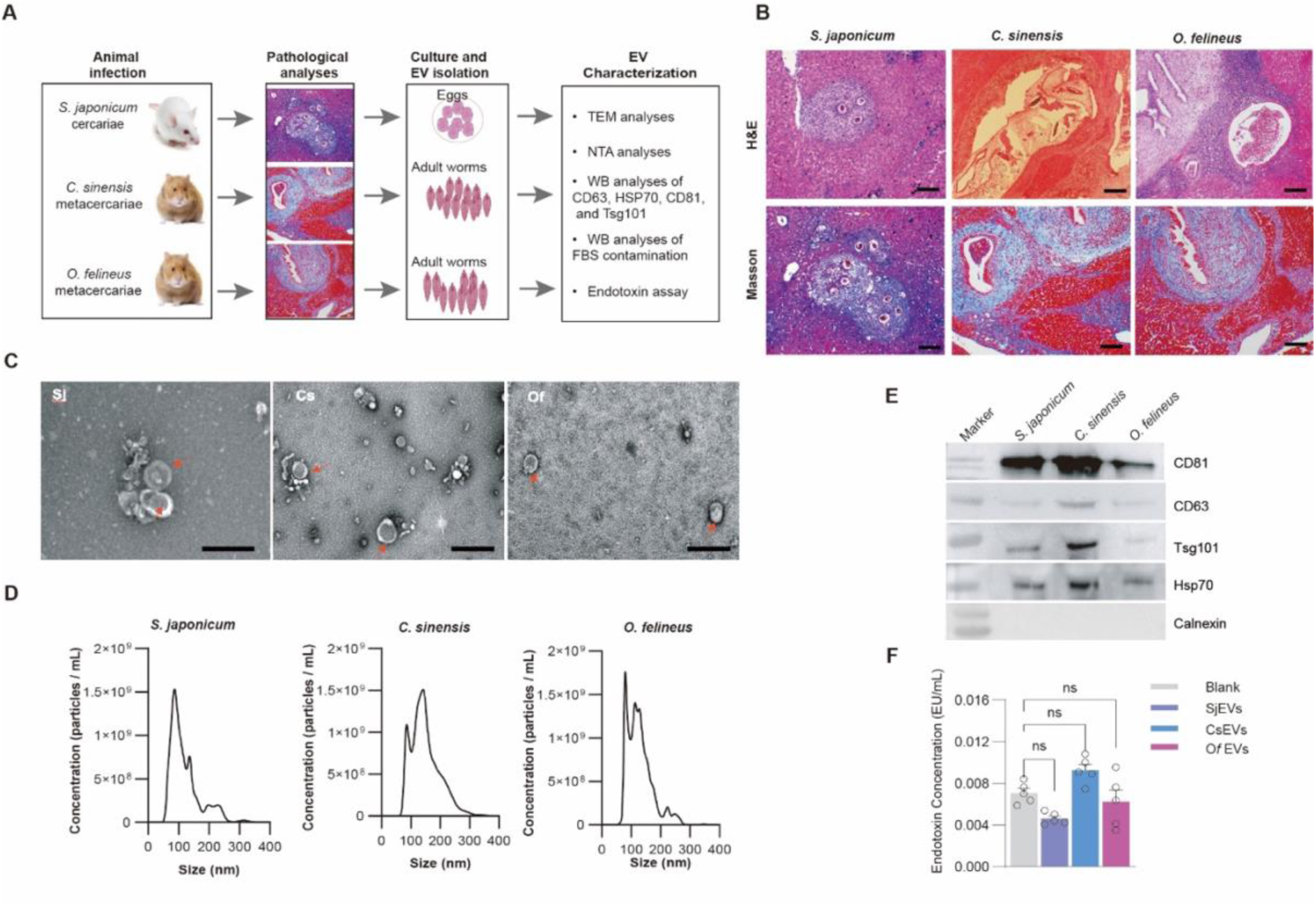
Characterization of isolated extracellular vesicles (EVs). (**A**) Schematic overview of EV isolation from *Schistosoma japonicum* (*S. japonicum*) eggs, adult *Clonorchis sinensis* (*C. sinensis*), and adult *Opisthorchis felineus* (*O. felineus*). (**B**) H&E and Masson staining analysis of liver fibrosis in the infected mice. Scale bars, 20 μm**. (C)** TEM analysis showing the morphology of isolated EVs. Red arrows indicate cup-shaped EVs. Scale bar = 500 nm. (**D**) Size distributions of isolated EVs. (**E**) Western blotting analysis of EV marker proteins including CD63, Tsg101, Hsp70, and CD81 as well as non-marker protein Calnexin in isolated EVs. **(F)** Endotoxin assay of isolated EVs. Data is presented as mean ± SEM. ns, not significant.

### 3.2 miRNome profiling of EVs from three trematodes

Small RNAs incorporated into EVs play a critical role in cell–cell communication ^35^. Determining the small RNA cargo in EVs can shed light on their regulatory functions. Therefore, we conducted de novo analyses to identify EV-associated small RNAs by RNA-seq. Bioinformatic analyses revealed that cis-regulatory small RNAs, tRNAs, rRNAs, and sRNAs constituted the dominant small RNA classes (**Figure 2A**). miRNAs were also considerably enriched in these parasite EVs (**Table S1**). More importantly, we found 18 conserved known miRNAs that were shared among all three EV populations, whereas *S. japonicum* harbored the highest number of species-specific miRNAs (n=40) compared with *O. felineus* EVs (n=9) (**Figure 2B**). In addition, we also identified 278 novel miRNAs in these parasite EVs (**Table S1**). Overall, these findings establish a comprehensive small RNome landscape of EVs across the three trematodes, highlighting a conserved core of miRNAs that may mediate cross-species host regulation.

**Fig 2.**
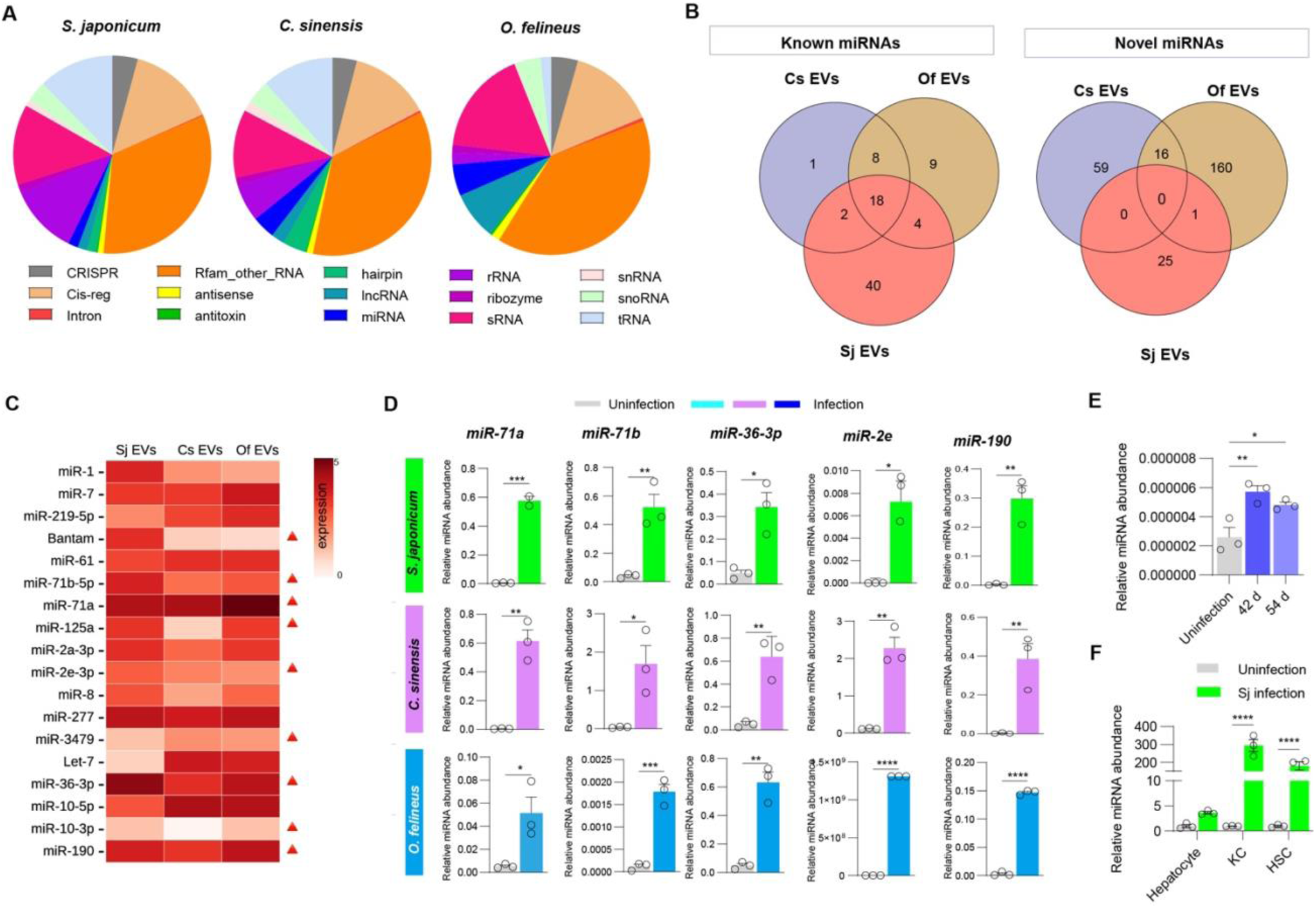
EV miRNome profiling from the key stages of three trematodes. **(A)** Pie charts showing the relative abundance of various small RNA categories (e.g., CRISPR, Cis-reg, Intron, rRNA, miRNA, etc.) in EVs from three trematodes. **(B)** Venn diagram displaying the overlap of known and novel miRNAs identified among the Cs EVs (blue), Of EVs (brown), and Sj EVs (red) groups. **(C)** Heatmap showing abundance of commonly known EV miRNAs across three trematodes (*S. japonicum*, *C. sinensis* and *O. felineus*). Red arrows indicate parasite specific miRNAs. **(D)** RT-qPCR validation of the abundance of selected common EV miRNAs in livers of infected mice (*S. japonicum*, *C. sinensis* and *O. felineus*). **(E)** RT-qPCR analyses of the abundance of *miR-36-3p* in livers of *S. japonicum* infected mice at different times. **(F)** RT-qPCR analyses of the abundance of *miR-36-3p* in isolated primary liver cells including hepatocytes, Kupffer cells, and hepatic stellate cells (HSCs) from infected mice at 42 dpi. Statistical analysis was conducted using Student’s *t*-test. Data in D-F are shown as mean ± SEM, * means *P* < 0.05, ** means *P* < 0.01, *** means *P* < 0.001, **** means *P* < 0.0001.

EV-mediated small RNAs could also play regulatory roles in a cross-species manner. RT-qPCR analysis of several abundant EV-derived miRNAs indicated their significant presence in the livers of infected animals (**Figure 2D**), suggesting that the EV miRNA cargo could be transferred into host liver cells. By focusing on the miRNAs shared among three parasites, we noted *miR-71a*, *miR-36-3p*, *miR-277*, and *miR-190* showed relatively high abundance in all species (**Figure 2C**). Moreover, *miR-71a*, *miR-36-3p*, and *miR-190* are likely parasite-specific miRNAs. Since *miR-71a* ^36^, *miR-2a* ^37^and miR-190 (manuscript submitted) were shown to suppress liver fibrosis in *S. japonicum* infected mice, we instead focused on *miR-36-3p*, whose functions have not been elucidated. First, we validated that this parasite-specific miRNA, *miR-36-3p*, was indeed enriched in the livers of infected mice (**Figure 2E**). Given that the liver is an organ composed of various cell types, we further determined which cell type accumulated *miR-36-3p*. The results indicated that *miR-36-3p* was significantly enriched in Kupffer cells (**Figure 2F**), suggesting that host Kupffer cells serve as the primary recipient cells for parasite-derived EVs.

### 3.3 Parasite EVs induced M1 polarization in Kupffer cells

Based on the above RT-qPCR results, parasite EVs were suggested to be mainly taken up by host Kupffer cells in the infected animal. Consequently, we first evaluated the effects of these EVs on Kupffer cell functions *in vitro*. We used PKH-26 to label *C. sinensis* EVs and the corresponding EV-depleted supernatant (EVDS) during EV isolation. RAW264.7 cells were treated with the labelled EVs or EVDS. The results indicated that parasite EVs can be taken up by recipient cells in a time-dependent manner (**Figure 3A**). Next, we evaluated whether EV miRNA cargos can be transferred into recipient cells. RT-qPCR analysis indicated that the abundance of selected EV miRNAs in recipient cells was significantly increased in the treated cells, indicating that parasite EV miRNA cargo can be transferred into the recipient cells (**Figure 3B**).

**Figure 3.**
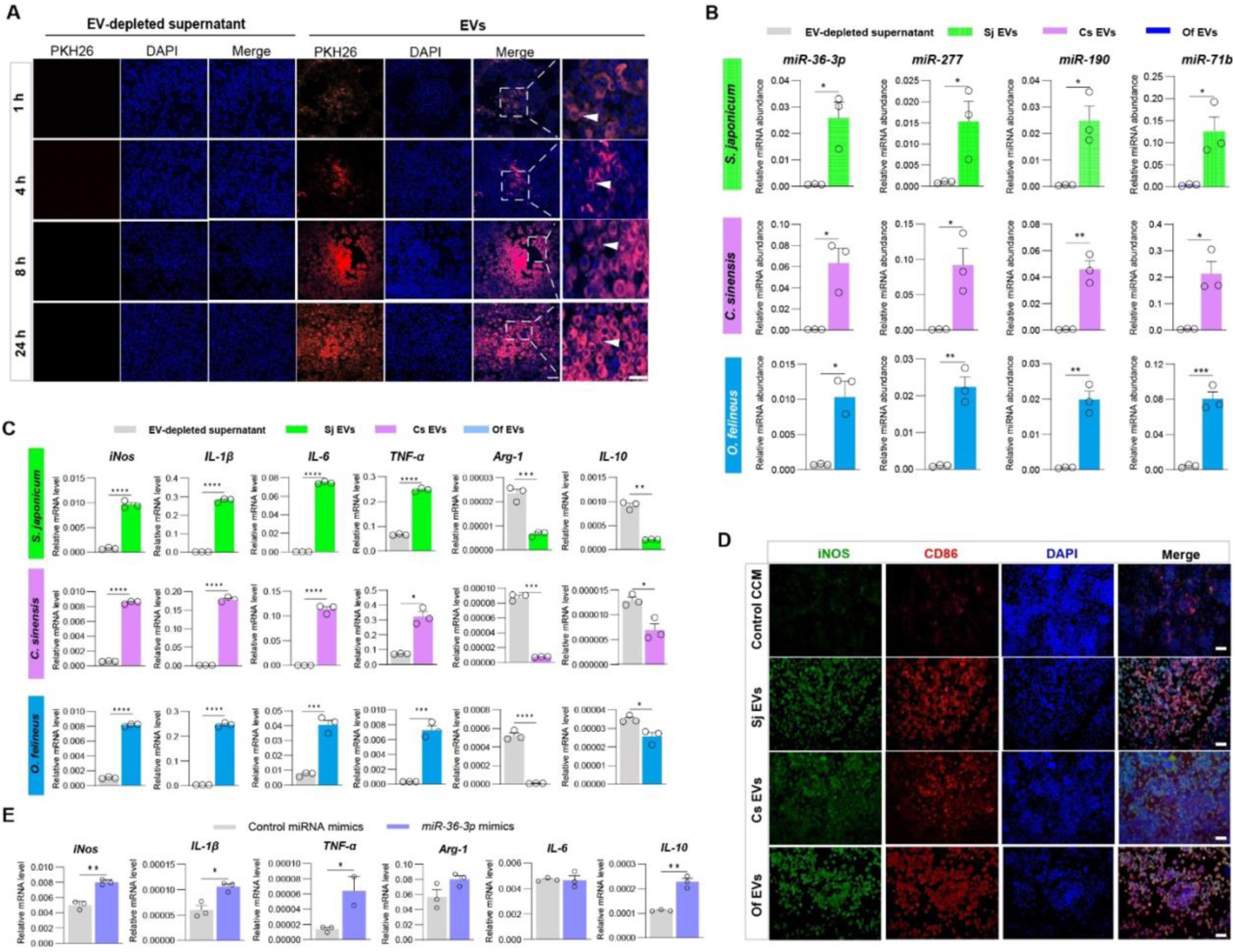
EVs are taken up by macrophages and influence their polarization. **(A)** Time-dependent internalization of PKH26-labelled EVs (red). PKH26-labelled EVs and EV-depleted supernatant (EVDS) were incubated with the macrophages at 1h, 4 h, 8 h, and 24 h. Bars represent 20 μm. **(B)** RT-qPCR analyses of the abundance of parasite EV miRNA in recipient cells treated with *S. japonicum* EVs, *C. sinensis* EVs, and *O. felinus* EVs, as well as their corresponding EV-depleted supernatant (EVDS).**(C)** RT-qPCR analysis of the transcript levels of M1/M2 marker genes in macrophages treated with *S. japonicum* egg EVs, *C. sinensis* EVs, and *O. felineus* EVs, as well as the corresponding controls (EV-depleted supernatant from each parasite, EVDS), respectively. **(D)** Immunofluorescence analysis of iNOS (green) and CD86 (red) expression in RAW264.7 cells exposed to *S. japonicum* EVs, *C. sinensis* EVs, and *O. felineus* EVs, as well as control culture-conditioned medium (CCM). Bars represent 20 μm. **(E)** RT-qPCR analysis of the transcript levels of M1/M2 marker genes in macrophages transfected with *miR-36-3p* mimics and control miRNA mimics. Data are shown as mean ± SEM from three independent experiments. \**P* means < 0.05, ** means *P* < 0.01, *** means *P* < 0.001, **** means *P* < 0.0001 (Student’s *t*-test)

Moreover, we determined the effects of parasite EVs on cell M1/M2 polarization. RT-qPCR results revealed that parasite EV-treated RAW264.7 cells underwent M1 polarization, as shown by elevated expressions of M1 markers (iNOS, CD86) and several cytokines (IL-1β, IL-6, TNF-α), whereas these increased expressions were not found in the cells treated with EVDS (**Figure 3C**). Consistently, immunofluorescence analysis validated that RAW264.7 cells treated with these parasite EVs showed high expression of iNOS and CD86 compared with the control culture-conditioned medium (CCM) (**Figure 3D**). In addition, the induced M1 polarization was alleviated when applying heat-inactivated parasite EVs (**Figure S2**), indicating that parasite EVs indeed play regulatory roles of cell functions. To determine whether the parasite EV miRNA cargo exerts a similar effect, we transfected *miR-36-3p* mimics into Kupffer cells (ImKCs). As expected, the cells underwent M1 polarization as determined by RT-qPCR. Taken together, these findings demonstrate that parasite EVs, and this EV miRNA cargo, at least miR-36-3p, reprogram Kupffer cells toward a pro-inflammatory M1 phenotype (**Figure 3E**).

### 3.4 Kupffer cells treated with parasite EVs or miR-36-3p cargo induced HSC activation

The interaction between Kupffer cells and hepatic stellate cells (HSCs) constitutes a key driver of liver fibrosis progression. To determine the effect of parasite EV-treated Kupffer cells on HSC activation, we first cultured RAW264.7 cells and treated them with parasite EVs or control condition culture medium (CCM). After thorough washing to remove the potential contamination from parasite EVs, HSCs (JS1) were co-cultured with the treated Kupffer cells (**Figure 4A)**. We found that cell viability and migration were increased upon parasite EV treatment. Notably, co-culture with the cells treated with parasite EVs led to HSC migration (**Figure 4B and 4C and Figure S3**). Further RT-qPCR analyses indicated that the transcript levels of *Acta2*, *Col1a1*, *Col3a1*, and *TGF-β* were significantly increased (**Figure 4D**). Next, we determined whether transfection of the EV miRNA cargo resulted in a similar effect. Expectedly, transfection with *miR-36-3p* significantly enhanced HSC viability and cell proliferation (**Figure 4E and 4F and Figure S4**) and HSC activation (**Figure 4G**) while co-transfection of *miR-36-3p* antisense mitigated these effects (**Figure 4F and 4G**).

**Fig 4.**
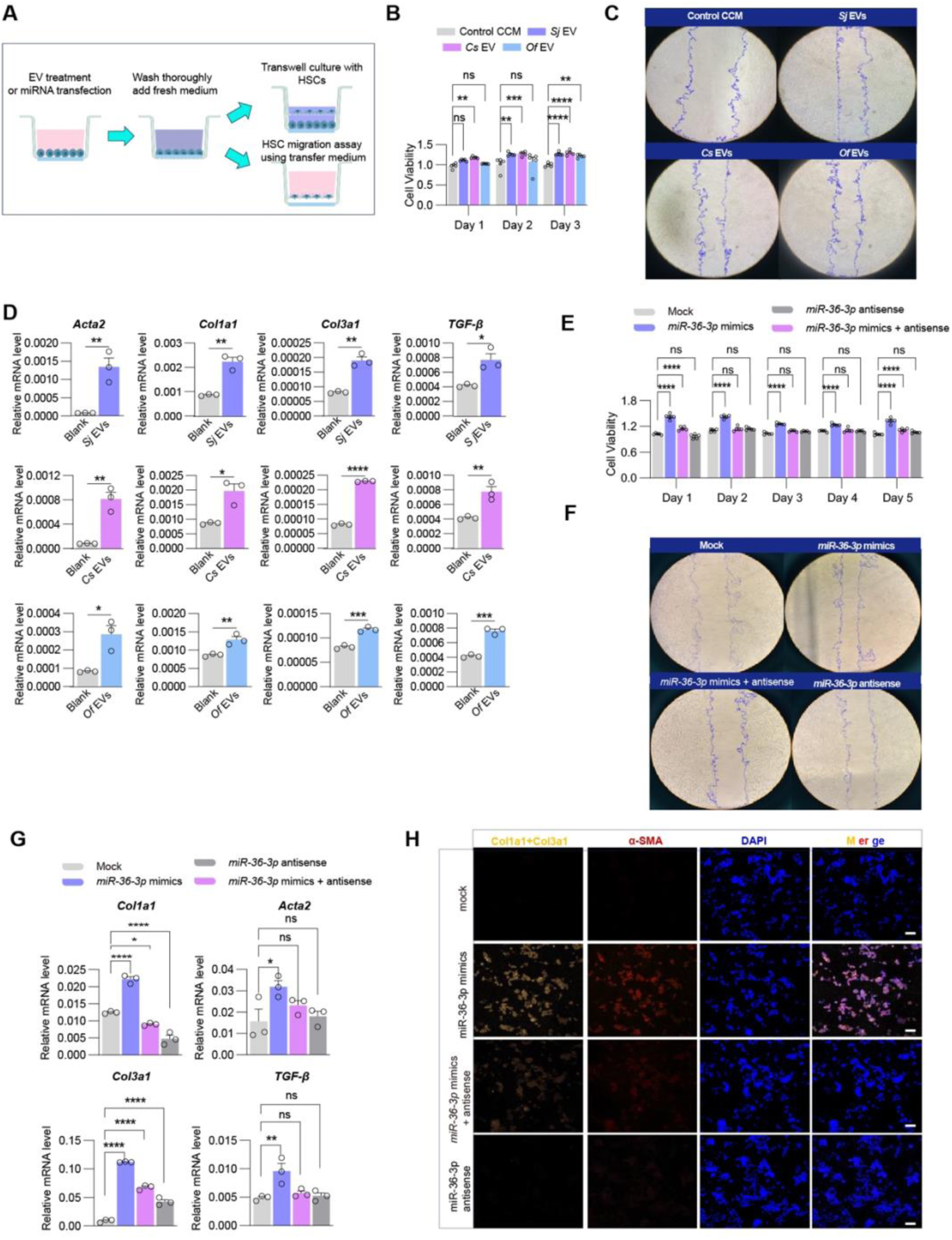
EVs and miRNA cargo regulate macrophage functions that drive HSC activation. **(A)** Schematic overview of cell culture strategies. **(B)** Cell viability of co-cultured HSCs with RAW264.7 cells treated with parasite EVs and CCM at day 1, 2, and 3 of post co-culture. **(C)** Representative results of HSC cell migration under co-culture conditions with RAW264.7 treated with parasite EVs and CCM. **(D)** RT-qPCR analysis of the transcript levels of *Col1a1, Col3a1*, *Acta2* and *TGF-β* in HSCs after co-culture with macrophage treated with different EVs and their corresponding EV-depleted supernatant (EVDS). **(E)** Cell viability of HSCs co-cultured with Kupffer cells transfected with *miR-36-3p* mimics and/or *miR-36-3p* antisense at 1-5 days of post-transfection. **(F)** Representative results of HSC cell migration after co-culture with Kupffer cells transfected with *miR-36-3p* mimics and/or *miR-36-3p* antisense. **(G)** RT-qPCR analysis of transcript levels of *Acta2, Col1a1, Col3a1,* and *TGF-β* in HSCs after co-culture with Kupffer cells transfected with *miR-36-3p* mimics and/or *miR-36-3p* antisense. **(H)** Immunofluorescence staining of Col3a1, Col1a1 and α-SMA in HSCs co-cultured with Kupffer cells transfected with *miR-36-3p* mimics and/or *miR-36-3p* inhibitor antisense. Scale bar = 20 μm. For panel B, D, F, and G, data are shown as mean ± SEM from three independent experiments. * means *P* < 0.05, ** means *P* < 0.01, *** means *P* < 0.001, **** means *P* < 0.0001 (one-way ANOVA or Student’s t-test was used for statistical analyses).

To corroborate these findings, we isolated primary Kupffer cells from murine livers, then transfected with *miR-36-3p* mimics and/or *miR-36-3p* antisense, and then co-cultured them with HSCs (JS1). Immunofluorescence analysis showed that transfection of *miR-36-3p* mimics into primary Kupffer cells indeed activated HSCs, whereas co-transfection with *miR-36-3p* antisense mitigated the effect (**Figure 4H and Figure S4**). Taken together, these results demonstrated that parasite EVs and their *miR-36-3p* cargo regulate HSC activation via Kupffer cells.

### 3.5 Identification of *miR-36-3p* targetome by single-cell RNA sequencing and bulk RNA-seq

To gain insight into the mechanism by which *miR-36-3p* mediated Kupffer cells to regulate HSC activation, we determined the targetome of *miR-36-3p* at the whole-organ level. To this end, we established an animal model by infecting mice with *S. japonicum* cercariae and then administering *miR-36-3p* antisense (**Figure 5A**). After four injections, the mice were sacrificed at 56 days post infection (dpi). We first used RT-qPCR to determine the abundance of the parasite-specific *miR-36-3p* in the livers of treated animals. The results indicated that the abundance of *miR-36-3p* was markedly reduced upon injection of *miR-36-3p* antisense (**Figure 5B**). In addition, pathological analyses using H&E, Masson, and immunohistochemistry staining (**Figure 5C**), followed by quantitative analysis (**Figure S5**), indicated that granuloma size and areas were significantly reduced in mice injected with *miR-36-3p* antisense (**Figure 5D**). Subsequently, RT-qPCR analysis showed that the transcript levels of *Col1a1*, *Col3a1*, *Acta2* and *TGF-β* in livers of mice treated with *miR-36-3p* antisense were significantly reduced (**Figure 5E**), indicating that liver fibrosis was mitigated. Additionally, the observation was further confirmed by measuring inflammation index in murine sera, such as AST, ALT, and liver HYP levels (**Figure 5F**). Overall, these results indicated that inhibiting parasite *miR-36-3p* alleviated liver fibrosis, validating this established animal model for identifying *miR-36-3p* targets.

**Fig 5.**
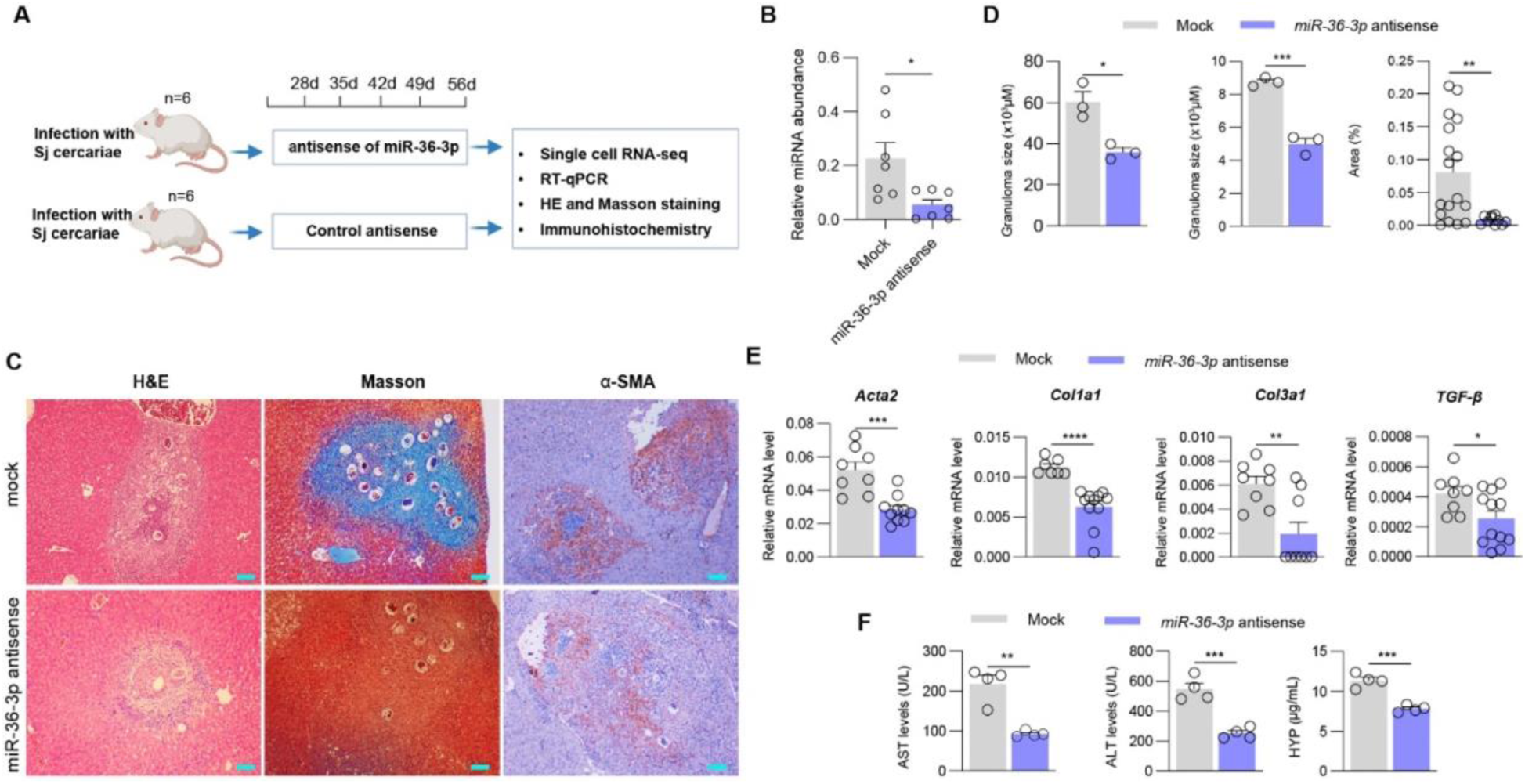
Establishment of an animal model for identifying *miR-36-3p* targetome. **(A)** Schematic illustration of the *S. japonicum* animal model for identifying *miR-36-3p* targetome. **(B)** RT-qPCR validation of the abundance of *miR-36-3p* in liver of mice administered *miR-36-3p* antisense. (**C**) Representative results of liver tissues analyzed by H&E, Masson staining, and immunohistochemistry assay for α-SMA. Scale bars =20 μm. (**D**) Quantitative analysis of granuloma size and collagen deposition area (%) based on H&E, Masson, and immunohistochemistry results. (**E**) RT-qPCR analysis of the mRNA levels of *Col1a1*, *Col3a1*, *Acta2* and *TGF-β* in the livers of mice administered *miR-36-3p* antisense. (**F**) Quantitative analysis of serum AST and ALT levels and liver HYP levels in mice administered *miR-36-3p* antisense. Data are presented as mean ± SEM. Statistical significance was determined using Student’s *t*-test. * means *P* < 0.05, ** means *P* < 0.01, *** means *P* < 0.001 (**B**, **D**, **E**, and **F**).

Therefore, we employed single-cell RNA-seq to globally map the *miR-36-3p* targetome. First, we isolated hepatic non-parenchymal cells (NPCs) from *miR-36-3p* antisense-treated and control mice, respectively, and performed scRNA-seq to determine altered gene expression. Following stringent filtering, a total of 10,112 high-quality cell transcriptomes were retained and resolved into 22 distinct clusters (**Figure 6A and 6B**). These clusters encompassed inflammatory monocytes, Kupffer cells, active macrophages, fibroblasts/HSCs, NK cells, mature/early B cells, plasma cells, activated/mature/immature neutrophils, and others. Compared with control liver cells, administration of *miR-36-3p* antisense resulted in 202 down-regulated and 344 up-regulated genes (**Figure 6C and 6F**). These differentially expressed genes were mainly enriched in Kupffer cells (n = 58), followed by inflammatory monocytes (n = 29), T cells (n = 29), mature macrophages (n = 27), and other cell types, suggesting that *miR-36-3p*-mediated regulation of liver fibrosis occurred predominantly in Kupffer cells.

**Figure 6.**
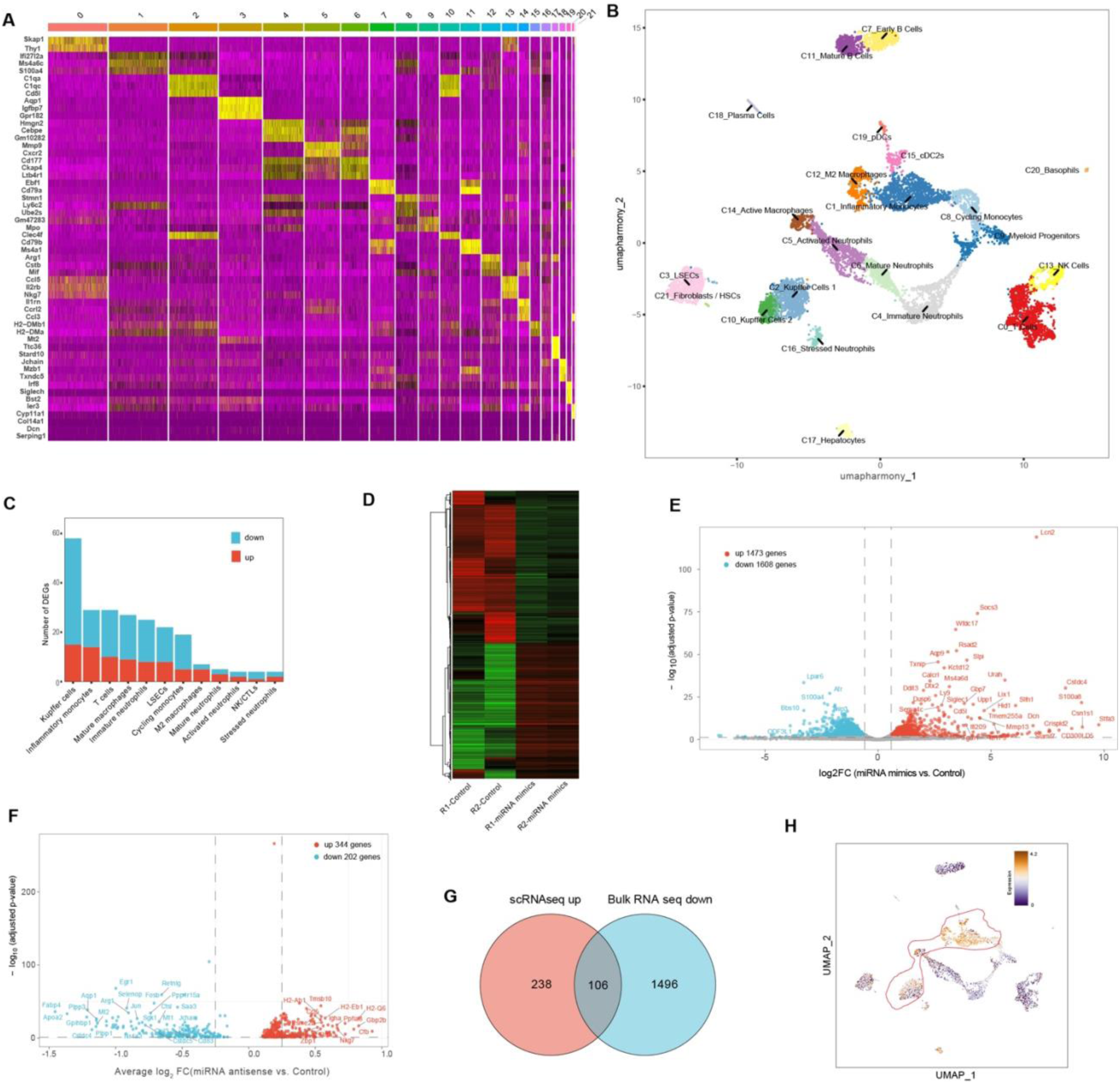
Identification of *miR-36-3p* targetome by single-cell RNA-seq and bulk RNA-seq. **(A)** Heatmap of marker gene expression across hepatic non-parenchymal cell subpopulations. **(B)** UMAP of different cell types based on single-cell RNA-seq data. **(C)** Number of differentially expressed genes (DEGs) in different cell types. **(D)** Heatmap showing DEGs from bulk RNA-seq analysis. R1 and R2 represent two biological replicates. **(E)** Volcano plot showing DEGs (bulk RNA-seq) in cells transfected *miR-36-3p* mimic as compared to control miRNA. **(F)** Volcano plot of global DEGs in mice administrated antisense of *miR-36-3p*. **(G)** Venn diagram illustrating the overlap between scRNA-seq upregulated genes (*miR-36-3p* antisense vs mock), bulk RNA-seq downregulated genes (*miR-36-3p* mimic vs mock). The numbers in the intersections represent the overlapping genes. **(H)** Single-cell expression profile of *Atox1* in hepatic cell subpopulations. Color intensity represents the expression level of the indicated gene, with red colors indicating higher expression.

Additionally, we further transfected cultured KCs with *miR-36-3p* mimics (**Figure 6D**) and performed bulk RNA-seq analysis on treated cells. Bioinformatic analyses indicated that there were 1,473 up-regulated and 1,608 down-regulated genes following *miR-36-3p* mimics treatment (adjusted *P* < 0.05, FC > 1.5; **Figure 6E**). To strengthen target identification, we focused on a total of 106 genes that were up-regulated in the single-cell RNA-seq data and simultaneously down-regulated in the bulk RNA-seq analyses. Next, we used RNAhybrid to analyze *miR-36-3p* target among these 106 genes (**Figure 6G**). Through this integrative approach, Atox1 emerged as a putative direct target of *miR-36-3p*. As expected, *Atox1* is also highly expressed in Kupffer cells according to the scRNA-seq data (**Figure 6H**). Overall, this integrative multi-omics screening identifies *Atox1* as a primary candidate target of *miR-36-3p*s.

### 3.6 *miR-36-3p* targeting of *Atox1* regulates HSC activation

Upon integration of targetome data from single-cell and bulk RNA-seq alongside stringent bioinformatics filtering, we identified *Atox1* as a candidate target of *miR-36-3p*. To validate this, we first assessed *Atox1* expression in cells transfected with *miR-36-3p* mimics using RT-qPCR. The results indicated that *Atox1* expression was significantly suppressed upon *miR-36-3p* mimics transfection, whereas co-transfection with *miR-36-3p* antisense reversed its expression (**Figure 7A**). We further validated that *in vivo* delivery of miR-36-3p antisense to mice significantly increased *Atox1* mRNA level compared with control-treated mice (**Figure 7B**). Next, we tested whether this regulatory effect occurred in cells treated with parasite EVs. The results showed that RAW264.7 cells treated with *S. japonicum* EVs (Sj EVs), *C. sinensis* EVs (Cs EVs), or *O. felineus* EVs (Of EVs) all exhibited decreased *Atox1* expression (**Figure 7C**). To further corroborate this regulatory effect, we used an animal model to assess *Atox1* expression in mice infected with *S. japonicum* or *C. sinensis*. The results showed that *Atox1* transcript levels in the liver were significantly decreased at different time points after parasite infection (**Figure 7D and 7E**). Furthermore, immunohistochemical analysis of liver sections from infected mice treated with *miR-36-3p* antisense consistently showed that *Atox1* was upregulated in antisense-treated mice compared with controls (**Figure 7G**). Next, we designed siRNAs to knock down *Atox1* expression. As shown in **Figure 7F**, the transcript level of *Atox1* was significantly reduced in cells transfected with siRNA. Transfection with *Atox1* siRNA decreased *Atox1* expression, similar to the effect of *miR-36-3p* mimics, and also induced a shift in macrophage polarization (**Figure S7**). Overall, these results confirm that *Atox1* is a direct functional target of *miR-36-3p*.

**Figure 7.**
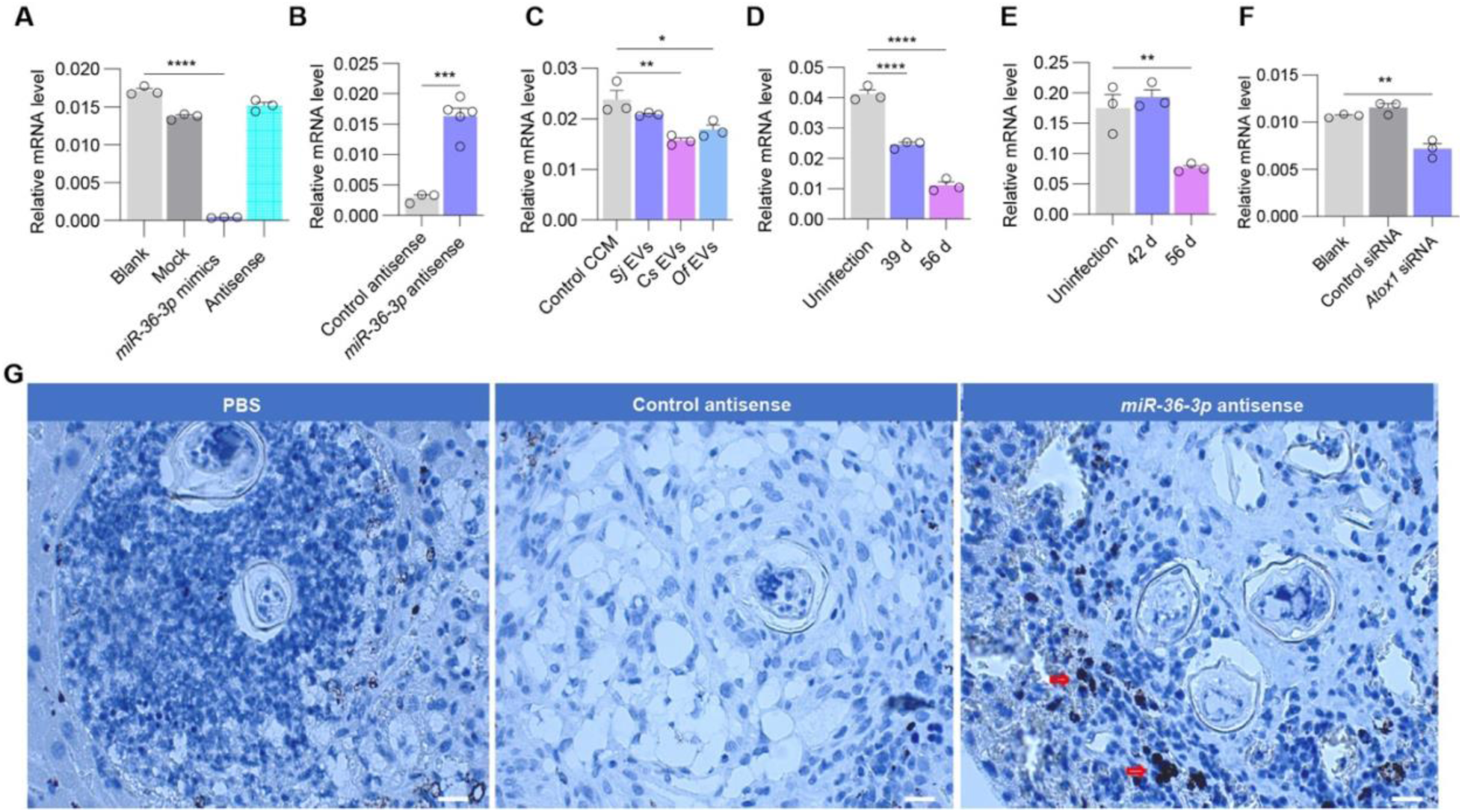
*Atox1* is a target of *miR-36-3p*. **(A)** RT-qPCR analysis of *Atox1* transcript levels in Kupffer cells that transfected with *miR-36-3p* mimics. Cells were treated with Lipofectamine (blank), control miRNA mimics (mock), *miR-36-3p* mimics, or *miR-36-3p* antisense. **(B)** RT-qPCR analysis of *Atox1* transcript levels in the livers of mice following *miR-36-3p* antisense injection. Mice were administered control antisense (mock) or *miR-36-3p* antisense. **(C)** RT-qPCR analysis of *Atox1* transcript levels in RAW264.7 cells treated with different parasite EVs and control CCM. **(D)** RT-qPCR analysis of hepatic *Atox1* transcript levels in mice infected with *S. japonicum* at various days. **(E)** RT-qPCR analysis of *Atox1* transcript levels in livers of infected mice with *C. sinensis* at various days. **(F)** RT-qPCR analysis of *Atox1* transcript levels in Kupffer cells transfected with siRNA. **(G)** Representative immunohistochemistry (IHC) images of Atox1 expression in mouse liver tissues from the antisense injection experiment. Red arrows indicate Atox1-positive cells. Scale bar =20 μm. Data are presented as mean ± SEM. One-way ANOVA (A, C, D-F) or Student’s *t*-test (B) was used for statistical comparisons. * means *P* < 0.05, ** means *P* < 0.01, *** means *P* < 0.001, **** means *P* < 0.0001.

### 3.7 The *miR-36-3p*–Atox1 axis regulates ROS levels in Kupffer cells

To reveal the regulatory mechanism of *miR-36-3p*-*Atox1* axis, STRING analysis identified a core protein–protein interaction (PPI) network of Atox1 including Cox17, Ccs, Slc31a1/2, Atp7a, Atp7b, and Sod3 (**Figure 8A**). KEGG analysis indicated that *Atox1* could regulate copper homeostasis and cellular redox status (**Figure 8A)**. Additionally, we noted that *Atox1* and its putative interactors were co-expressed at relatively high levels in inflammatory monocytes and Kupffer cells based on our scRNA-seq data, suggesting their potential interaction in regulating specific cell functions (**Figure 8B**). Furthermore, we assessed the expression of putative ATOX1 interactors in *Atox1* knock down (KD) cells using the *Atox1* siRNA. The results indicated marked downregulation of *Atp7a*, *Atp7b*, and *Ccs* (**Figure 8D**). *Atox1* KD by either *Atox1* siRNA or *miR-36-3p* mimics significantly increased intracellular copper levels in the treated cells (**Figure 8E**), reflecting its regulatory role in copper homeostasis. Moreover, we measured intracellular ROS levels and the results indicated that ROS was significantly elevated in both *Atox1* siRNA- and *miR-36-3p* mimics treated cells (**Figure 8C**), revealing a link between copper ions and redox homeostasis. Overall, these findings indicated that *Atox1* can interactively regulate the expression of copper transport–related genes to maintain ROS homeostasis in Kupffer cells.

**Figure 8.**
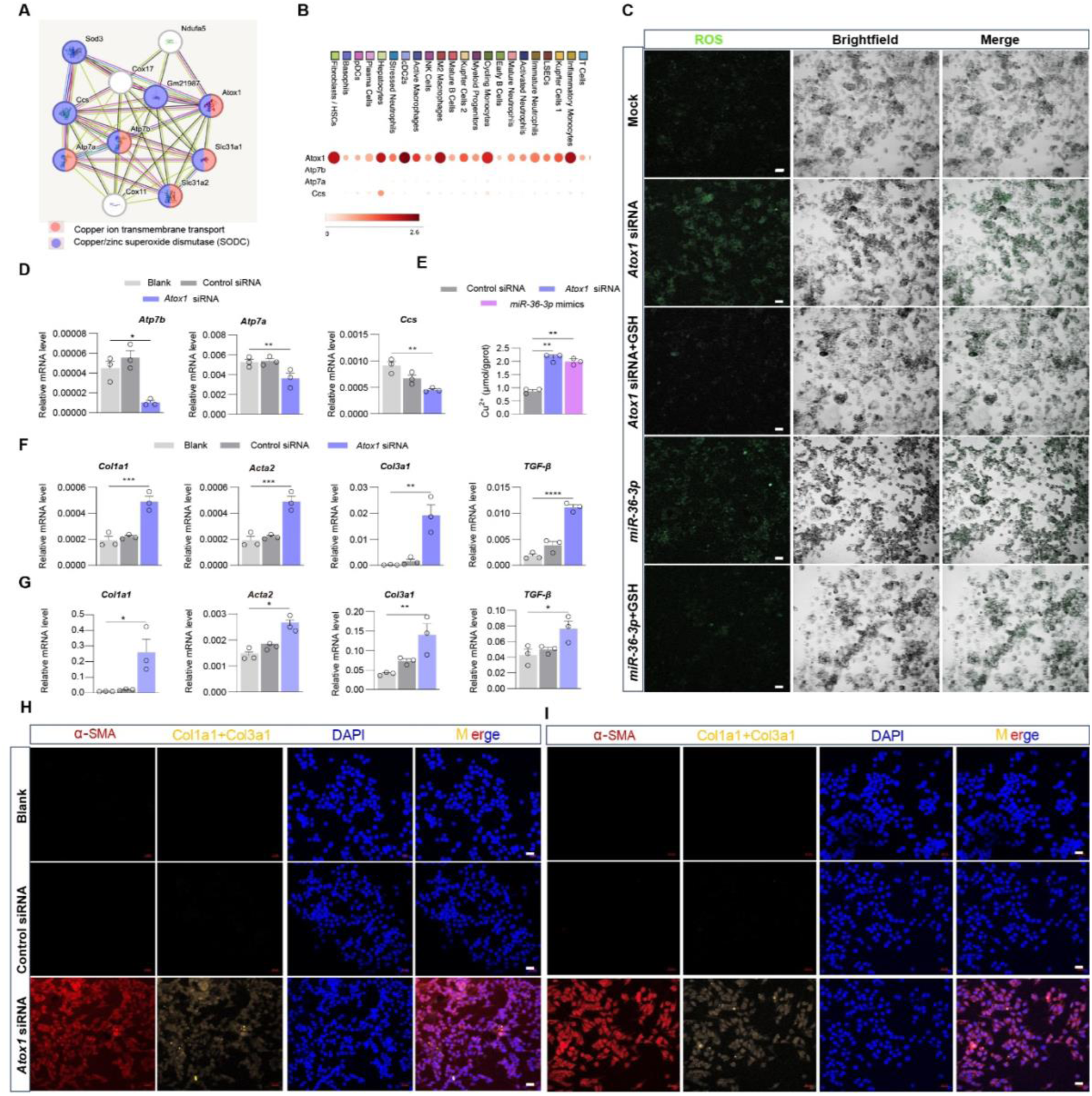
Suppression of Atox1 decreases cell proliferation and HSC activation. **(A)** STRING analysis of the Atox1-mediated protein–protein interaction network. **(B)** Single-cell RNA-seq (scRNA-seq) data showing the transcript levels of these interactive genes within Kupffer cells. **(C)** *Atox1* knockdown (KD) by either siRNA or *miR-36-3p* mimics leads to increased ROS in Kupffer cells. Scale bar = 20 μm. **(D)** RT-qPCR analysis of the transcript levels of selected putative Atox1 interactors in cells transfected with *Atox1* siRNA. **(E)** *Atox1* KD leads to intracellular copper accumulation in Kupffer cells. KCs were transfected with control, *Atox1* siRNA, or *miR-36-3p* mimics, and intracellular copper content was measured by a colorimetric assay using a copper-specific probe. **(F)** *Atox1* KD in Kupffer cells resulted in increased expression of pro-fibrotic genes in co-cultured HSCs (JS1). **(G)** *Atox1* KD in primary Kupffer cells resulted in increased expression of pro-fibrotic genes in co-cultured HSCs (JS1). **(H)** Immunofluorescence analysis of Col1a1, α-SMA, and Col3a1 expression in HSCs co-cultured with *Atox1* KD Kupffer cells. **(I)** Immunofluorescence analysis of Col1a1, α-SMA, and Col3a1 expression in HSCs co-cultured with *Atox1* KD primary Kupffer cells. Scale bar =20 μm. Data in **D**, **E**, **F**, and **G** are presented as mean ± SEM. Statistical analysis was conducted using one-way ANOVA. * means *P* < 0.05, ** means *P* < 0.01, ***means *P* < 0.001, **** means *P* < 0.0001.

Since transfection of *miR-36-3p* mimics into Kupffer cells resulted in activation of co-cultured HSCs, accompanied by a decrease in *Atox1* expression, we next determined whether *Atox1* knockdown (KD) by siRNA influenced HSC activation. Kupffer cells were transfected with *Atox1* siRNA and subsequently co-cultured with HSCs (JS1). As expected, *Atox1* knockdown activated co-cultured HSCs (**Figure 8F and 8H**) and induced a shift toward M1 polarization in Kupffer cells (**Figure S6**). To further corroborate these results, we isolated primary Kupffer cells from murine livers and treated them with *Atox1* siRNA. Consistently, *Atox1* knockdown in primary Kupffer cells activated co-cultured HSCs, as determined by RT-qPCR (**Figure 8G**) and immunofluorescence staining (**Figure 8I**). Taken together, these results demonstrate that *Atox1* KD in Kupffer cells promotes HSC activation, revealing the regulatory role of *Atox1* in hepatic fibrogenesis.

### 3.8 AAV sponge-mediated suppression of *miR-36-3p* alleviates liver fibrosis in mice infected with all three trematode species

To determine whether suppression of parasite *miR-36-3p* attenuates liver fibrosis in animal models, we designed a *miR-36-3p* sponge and packaged it into recombinant AAV. On day 10 post-infection, the animals were administered AAV-miR-36-3p sponge (anti-*miR-36-3p* AAV sponge), control AAV, or PBS via the tail vein. At 42 days post-infection, the animals were euthanized and the livers were collected for subsequent analyses (**Figure 9A**). First, we confirmed that parasite-specific *miR-36-3p* levels were effectively decreased in infected animals following AAV sponge injection, as verified by RT-qPCR (**Figure 9B**). As expected, immunohistochemical analysis showed that ATOX1 expression was increased in the livers of infected mice after AAV-*miR-36-3p* sponge injection (**Figure 9C**). RT-qPCR analysis indicated decreased expression of *Col1a1*, *Acta2*, and *Col3a1* in the livers of mice treated with AAV-*miR-36-3p* sponge compared with controls (**Figure 9D**). Additionally, histochemical and Masson staining analyses, together with quantitative analyses using ImageJ, revealed decreased granuloma size and reduced collagen accumulation (**Figure 9E and 9F**; **Figure S7**), indicating that liver fibrosis had been alleviated by blockade of the parasite *miR-36-3p*. These observations were further corroborated by measuring serum AST and ALT as markers of hepatic injury and HYP as a marker of collagen deposition (**Figure S8**). Overall, these findings provide in vivo proof-of-concept that therapeutic blockade of conserved parasite EV miR-36-3p restores *Atox1* expression and effectively attenuates liver fibrosis across all three trematode species.

**Figure 9.**
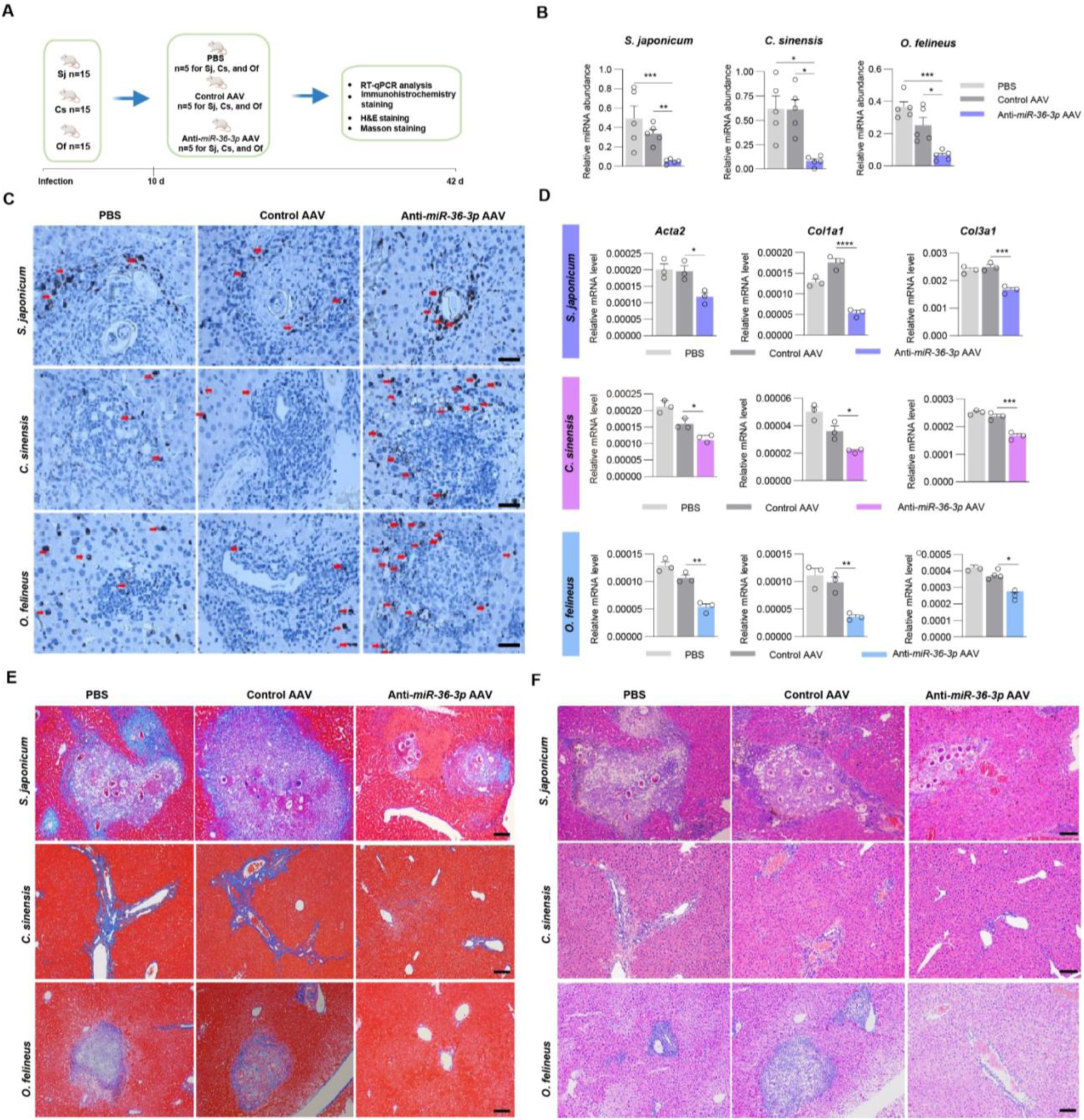
Suppression of *miR-36-3p* by AAV sponge alleviates liver fibrosis in mice infected with three trematodes. **(A)** Schematic diagram of animal experiment for AAV sponge injection. **(B)** Assessment of *miR-36-3p* abundance in the livers of infected mice administered with AAV sponge. **(C)** Immunohistochemistry analysis of Atox1 expression in liver sections from the AAV sponge administered animals. Red arrows indicate Atox1-positive cells. Scale bar = 20 μm. **(D)** RT-qPCR analysis of fibrosis-related gene expression in the livers of infected mice treated with AAV sponge. **(E)** Masson staining analysis of liver sections from mice administered with AAV sponge. Scale bar =20 μm. **(F)** H&E staining analysis of liver tissues from different treatments. Scale bar = 20 μm. Data in B and D are presented as mean ± SEM. One-way ANOVA was used for statistical analyses. * means *P* < 0.05, ** means *P* < 0.01, ***means *P* < 0.001, **** means *P* < 0.0001.

## 4 Discussion

Liver fibrosis caused by trematodes such as *S. japonicum*, *C. sinensis*, and *O. felineus* remains a significant global public health problem ^1, 38, 39^. Extracellular vesicles act as vital pathological agents in a cross-species communication manner and play a pivotal role in host–parasite interactions ^40^, yet the conserved mechanisms by which these trematode EVs promote liver fibrosis remain poorly defined. In this study, we successfully isolated and characterized EVs from three zoonotic trematodes, identified their EV miRNA repertoires for key etiological stages, and found that *miR-36-3p* is a shared EV miRNA cargo among these trematodes. In terms of functionality, *miR-36-3p* promoted M1-like polarization of Kupffer cells and subsequently facilitated HSC activation and liver fibrosis. Mechanistically, we used single-cell RNA sequencing from the whole organ level and bulk RNA seq from cell levels to comprehensively characterize the targetome of *miR-36-3p*. Upon stringent selection, we identified the copper chaperone *Atox1* as a direct target of *miR-36-3p*, whose suppression elevates intracellular copper and ROS levels to drive the activation of HSC. Importantly, our findings reveal a conserved parasite EVs miRNA cargo, *miR-36-3p*, that mediated host immune cells (Kupffer) by regulating copper/ROS homeostasis to promote HSC-driven fibrogenesis. More importantly, administration of AAV8 engineered *miR-36-3p* sponge in each trematode-infected animal model significantly halted liver fibrosis. Taken together, our findings revealed this common EV miRNA cargo of zoonotic trematodes could be a promising target for inhibiting helminth-induced liver fibrosis.

Extracellular vesicles carry diverse cargoes, including proteins, lipids, and miRNAs ^41, 42, 43^. In particular, EV miRNA cargo has been shown to play regulatory roles in pathological response of host in these trematodes. For instance, in *S. japonicum* eggs, their EVs contain *miR-33* and *miR-71a* that can regulate liver fibrosis ^12, 13, 36^. In *C. sinensis*, worm EVs were shown to promote the malignant phenotypes of cholangiocarcinoma and miRNA cargo such as *Csi-let-7a-5p* can activate pro-inflammatory macrophages to induce biliary injury^14^. In *O. felineus*, parasite can secrete exosome-like vesicles that can be uptaken by human cholangiocytes ^9^. However, whether there are shared miRNAs that exert conserved regulatory functions across these trematodes, resulting in similar clinical phenotype, remains a critical gap in the field.

Our work provides a comprehensive comparative landscape of EVs from these three important trematodes, particularly their miRNA cargo, by identifying the repertoire of conserved miRNAs. We revealed that *miR-36-3p* was present in all three parasite species and could be delivered into host cells, highlighting its conserved role in parasite–host communication. Next, we identified that *miR-36-3p* can target *Atox1,* which is a copper chaperone for maintaining intracellular copper homeostasis and redox balance ^44^. Accumulating evidence has demonstrated that copper is selectively delivered to distinct intracellular destinations via copper chaperones, such as *Atox1*, *Atp7a*, *Atp7b*, and *Ccs*, which delivers copper to *Sod1* ^45, 46, 47, 48^. This notion is further supported by our observation that knockdown of *Atox1* resulted in concomitant changes in both intracellular copper content and the reduced expression of related genes. Given the essential role of copper chaperones in maintaining cellular copper balance, we provide evidence linking Atox1-mediated copper homeostasis to the progression of liver fibrosis induced by parasitic infection, disruption of this homeostatic network inevitably elevates oxidative burden ^49^ and fosters an inflammatory milieu within cells ^50^.

Taken together, our findings demonstrate that an EV miRNA cargo, *miR-36-3p*, commonly shared by three trematodes, drives hepatic fibrogenesis. Mechanistically, *miR-36-3p* target *Atox1*, disrupting intracellular copper balance and leading to oxidative stress and inflammatory responses that activate HSCs, ultimately culminating in extracellular matrix deposition. Our findings uncover a previously unrecognized cross-species mechanism of parasite–host interaction and identify EV *miR-36-3p* cargo as a potentially promising therapeutic target for halting liver fibrosis caused by these trematodes.

## 5 Conclusion

In this study, we characterized EVs from three clinically important trematodes—*Schistosoma japonicum*, *Clonorchis sinensis*, and *Opisthorchis felineus*—and identified a conserved EV-derived miRNA repertoire shared across all three species. Among these, *miR-36-3p* directly targets *Atox1* in Kupffer cells, driving copper and ROS accumulation and subsequently activating hepatic stellate cells. In vivo delivery of an AAV8-encoded *miR-36-3p* sponge markedly attenuated liver fibrosis in all three infection models. These findings establish a conserved, parasite-derived EV miRNA cargo as a previously unrecognized driver of host liver pathology and a promising therapeutic target for parasite-induced liver fibrosis.

## Author Contributions

Lu Liu: conceptualization, methodology, formal analysis, investigation, data curation, Writing – original draft, visualization. Zhigang Lu: methodology, formal analysis, data curation, writing – review and editing, visualization. Shiyu Zhu, Chuantao Fang, Xiaoli Yan, Chunren Wang, Feng Liu, and Yanyan Zhang: methodology, resource. Maria Pakharukova: conceptualization, resource, writing – review and editing. Guofeng Cheng: conceptualization, resource, writing – review and editing, supervision, funding acquisition.

## Acknowledgements

This study was, in part or in whole, supported by National Natural Science Foundation of China (Grants No. 32361133554 to GC). The study is also sponsored by Shanghai Tongji University Education Development Foundation.

## Data Availability Statement

The data that support the findings of this study are available from the corresponding author upon reasonable request.

## Supplementary information

### Supplementary Figures

**Figure S1** Western blotting analysis of parasite EVs using specific antibodies against FBS.

**Figure S2.** RT-qPCR analysis of the transcript levels of M1/M2 marker genes in macrophages treated with *C. sinensis* EVs, boiled EVs and EV-depleted supernatant.

**Figure S3** Quantitative analysis of the images of HSC migration at the co-culture conditions with RAW264.7 cells.

**Figure S4** Quantitative analysis of the images of HSC migration at the co-culture conditions with Kupffer cells.

**Figure S5** Quantitative analysis of liver fibrosis parameters. Granuloma size was analyzed using ImageJ. Data are presented as mean ± SEM. \*\*\**P* < 0.001.

**Figure S6** *Atox1* KD induced a shift toward M1 polarization in Kupffer cells.

**Figure S7** Quantitative analysis of liver fibrosis parameters in mice treated with AAV *miR-36-3p* sponge. (A) Quantification of histological changes assessed by HE staining. (B) Quantification of fibrosis assessed by Masson staining. Data are presented as mean ± SEM. *P < 0.05, **P < 0.01, ***P < 0.001, ****P < 0.0001.

**Figure S8** Measurement of the biomarkers of hepatic injury in mice treated with AAV miR-36-3p sponge.

### Supplementary Tables

**Table S1.** The list of miRNAs identified in EVs isolated from three trematodes.

**Table S2.** The list of DEGs identified by single cell RNA seq.

**Table S3.** The list of DEGs identified by Bulk RNA seq.

**Table S4.** The list of primers and siRNA used in the present study.

