## Supplementary Fig S1-S8 for "Comparative landscape of extracellular vesicles across three pathogenic trematodes identifies a key conserved miRNA cargo involved in host liver fibrosis"

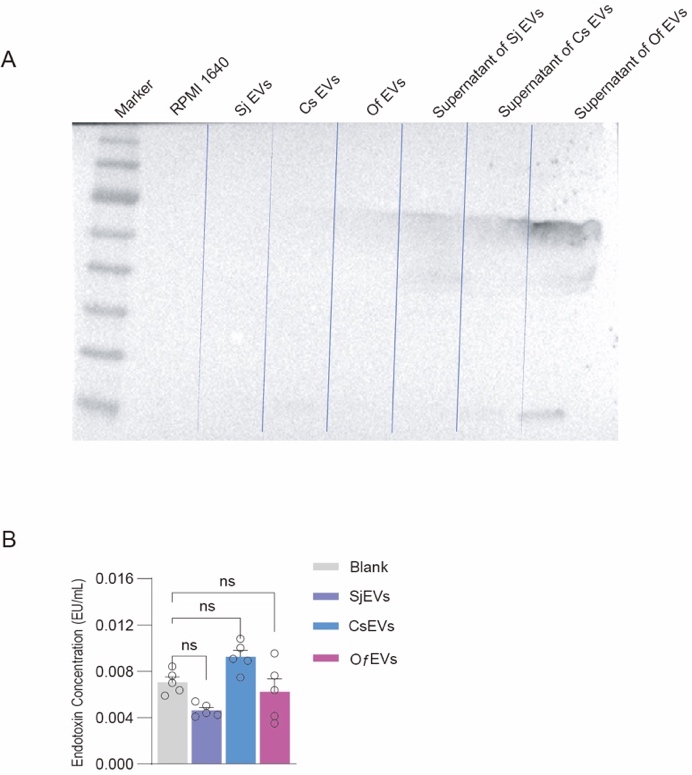

**Figure S1** Western blotting analysis of parasite EVs using specific antibodies against FBS.

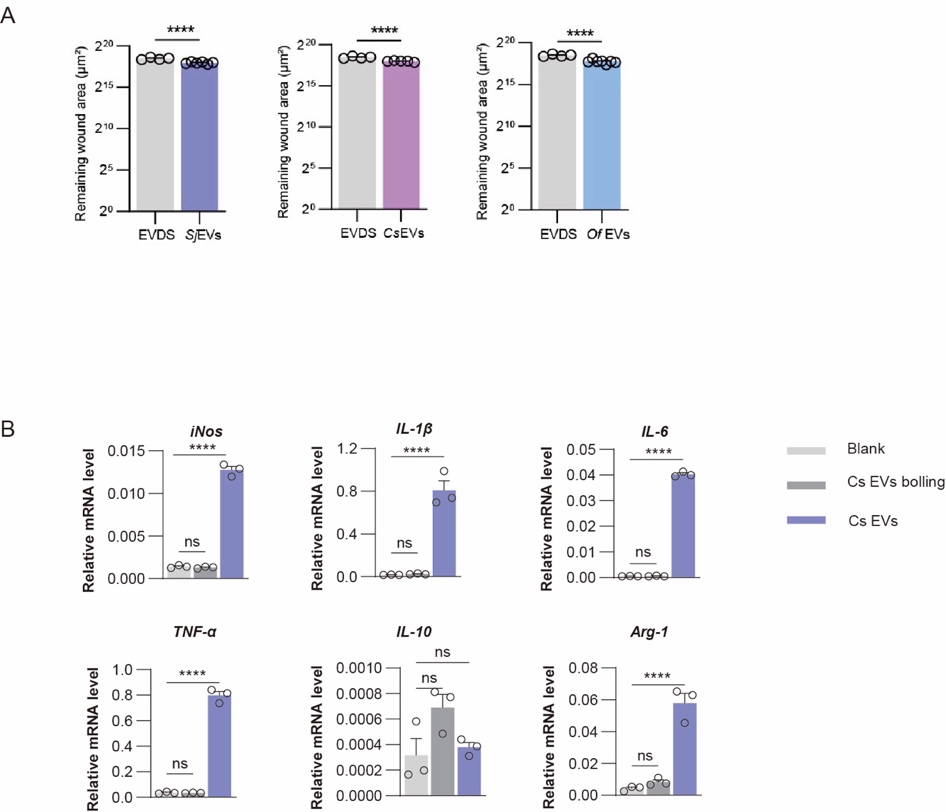

**Figure S2** RT-qPCR analysis of the transcript levels of M1/M2 marker genes in macrophages treated with *C. sinensis* EVs, boiled EVs and EV-depleted supernatant. Data is presented as mean ± SEM. *****P* < 0.0001.

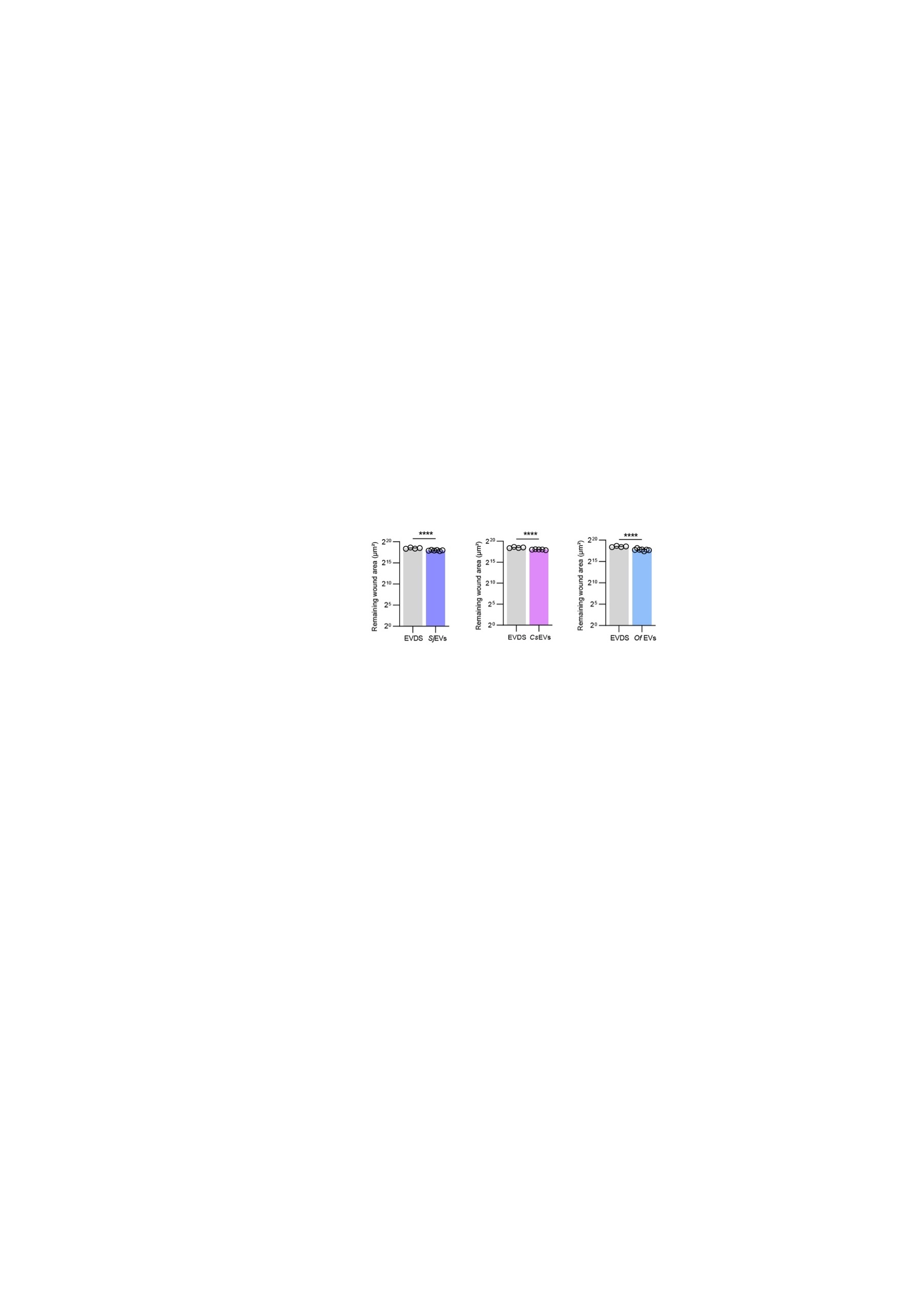

**Figure S3** Quantitative analysis of the images of HSC migration at the co-culture conditions with Raw264.7 cells. Data is presented as mean ± SEM. *****P* < 0.0001.

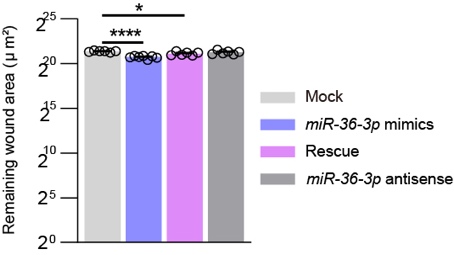

**Figure S4** Quantitative analysis of the images of HSC migration at the co-culture conditions with Kupffer cells. Data is presented as mean ± SEM. **P* < 0.05, *****P* < 0.0001.

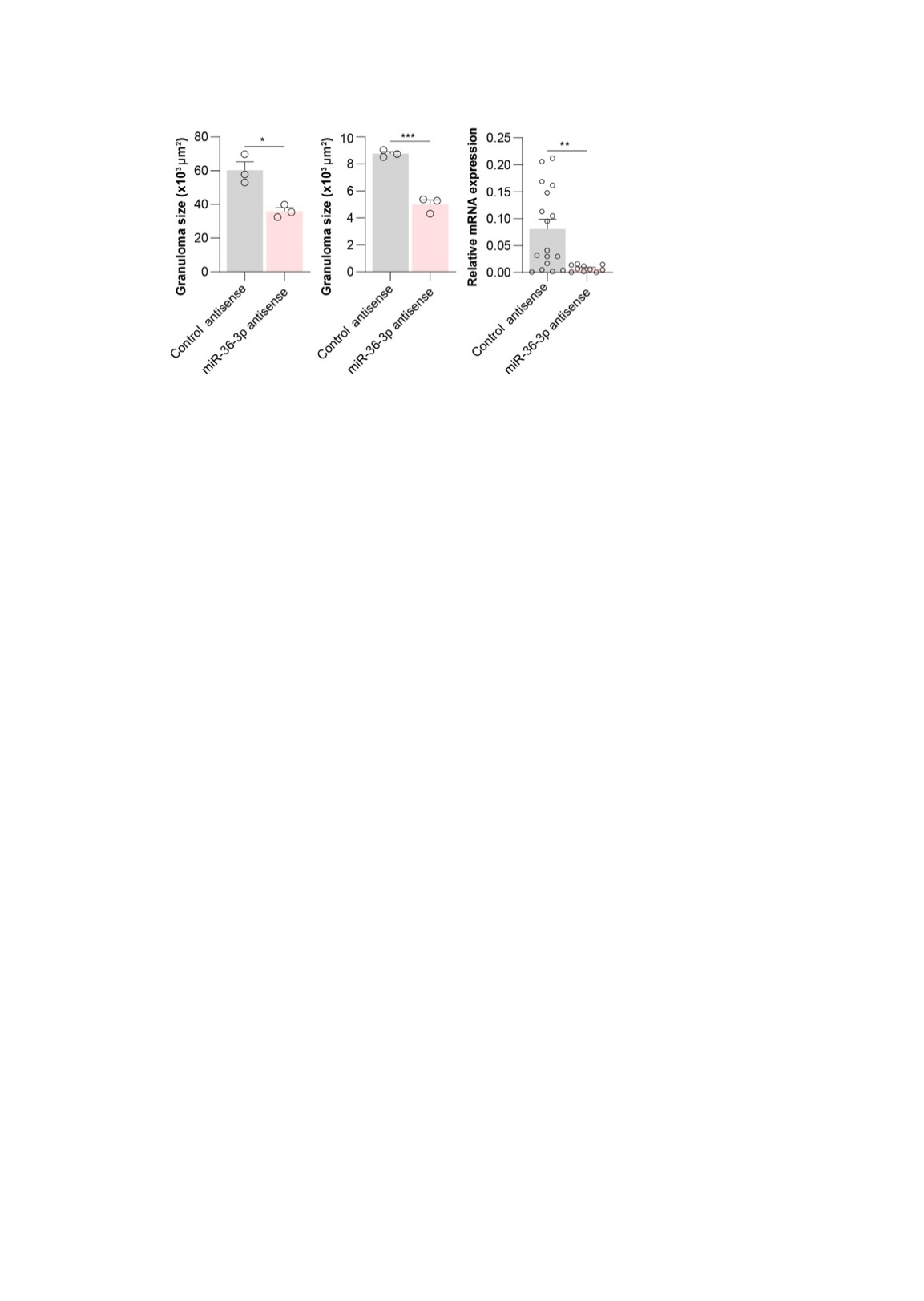

**Figure S5** Quantitative analysis of liver fibrosis parameters. Granuloma size was analyzed using ImageJ. Data are presented as mean ± SEM. **P* < 0.05, ***P* < 0.01, ****P* < 0.001.

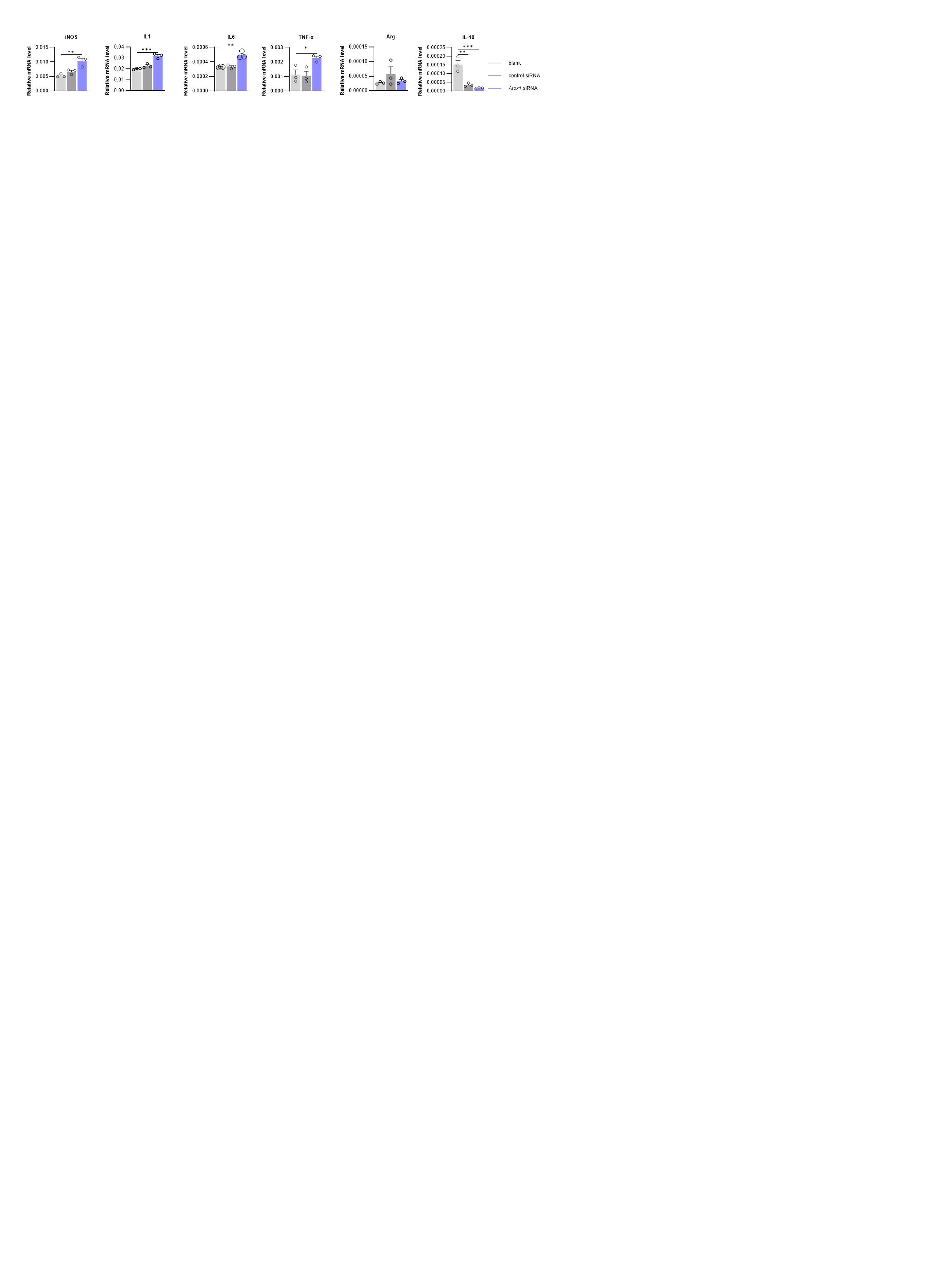

**Figure S6** *Atox1* KD induced a shift toward M1 polarization in Kupffer cells. Data is presented as mean ± SEM. ***P* < 0.01, ****P* < 0.001.

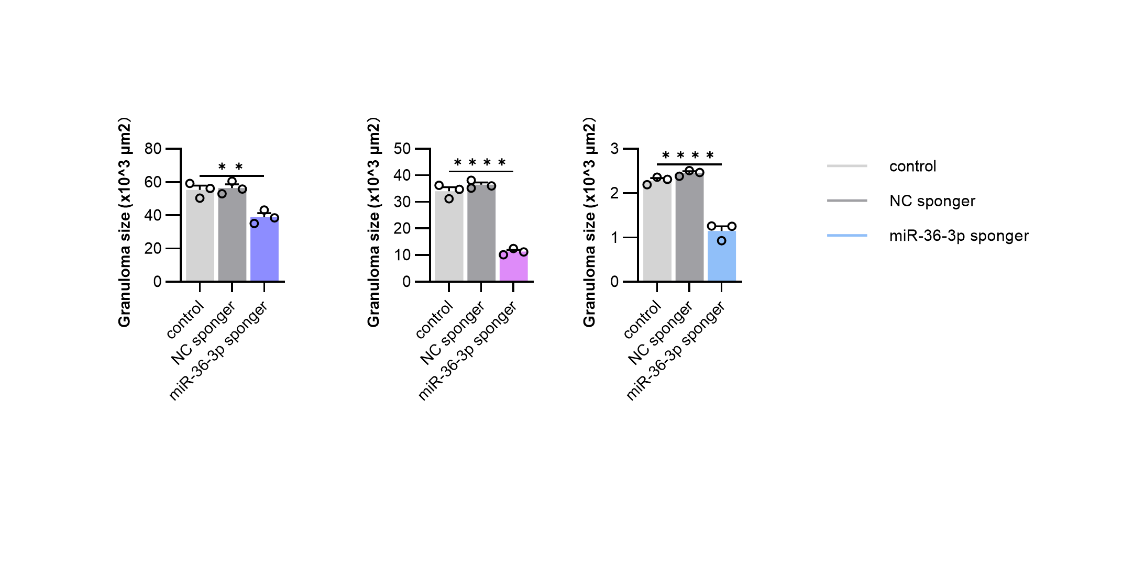
A

B

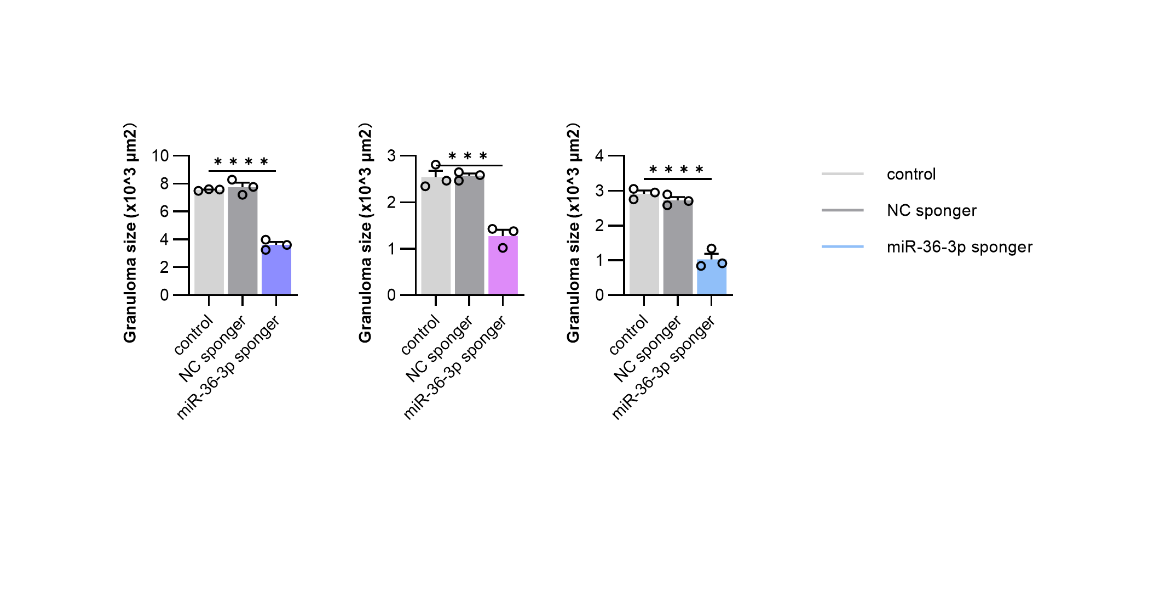

**Figure S7** Quantitative analysis of liver fibrosis parameters in mice treated with AAV *miR-36-3p* sponge. Granuloma size was analyzed using ImageJ. (A) Quantification of histological changes assessed by HE staining. (B) Quantification of fibrosis assessed by Masson staining. Data are presented as mean ± SEM. ***P* < 0.01, ****P* < 0.001, *****P* < 0.0001.

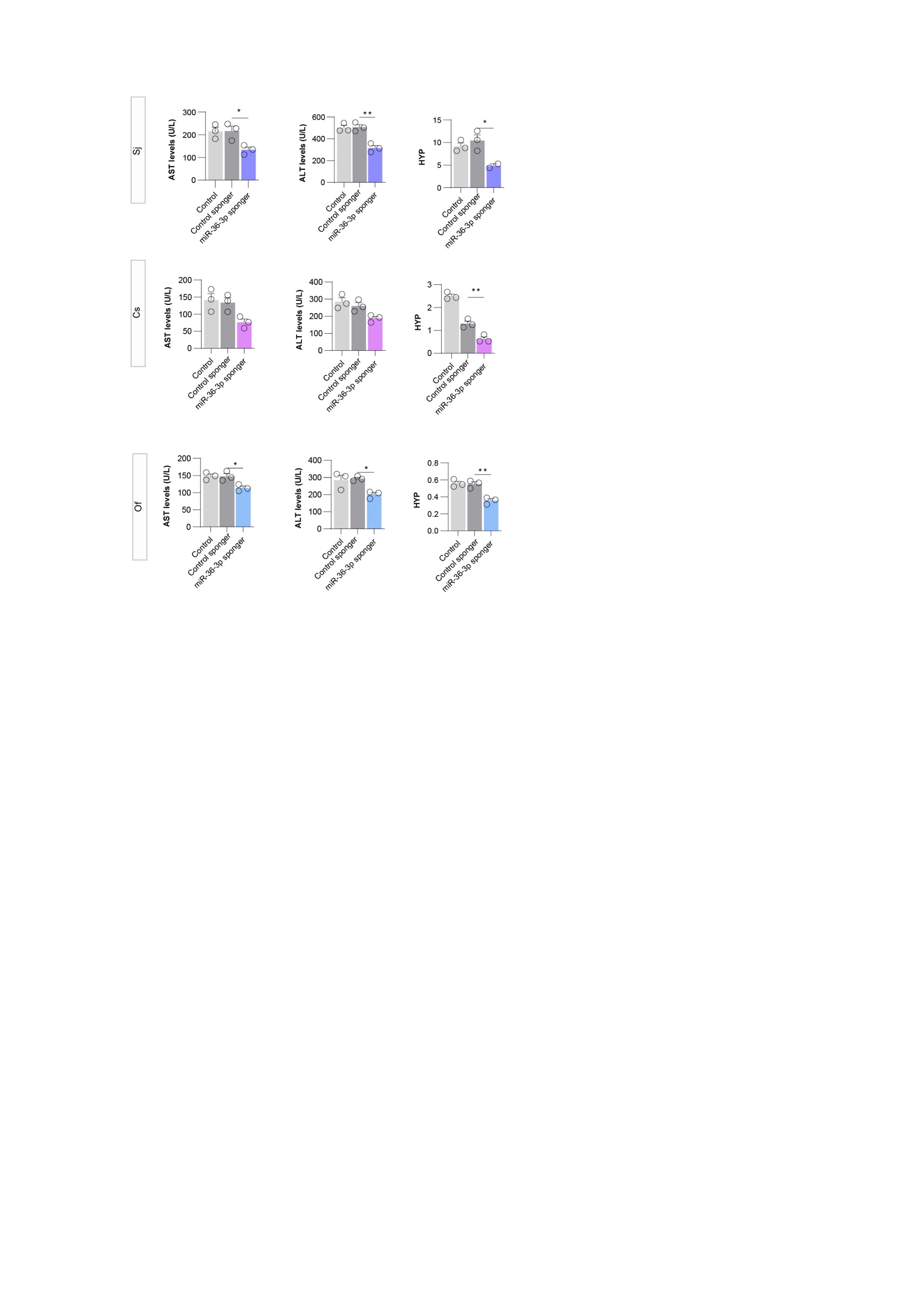

**Figure S8** Measurement of the biomarkers of hepatic injury in mice treated with AAV miR-36-3p sponge. Data is presented as mean ± SEM. **P* < 0.05, ***P* < 0.01.
.
